# A Statistical Framework for QTL Hotspot Detection

**DOI:** 10.1101/2020.08.13.249342

**Authors:** Po-Ya Wu, Man-Hsia Yang, Chen-Hung Kao

## Abstract

Quantitative trait loci (QTL) hotspots (genomic locations enriched in QTL) are a common and notable feature when collecting many QTL for various traits in many areas of biological studies. The QTL hotspots are important and attractive since they are highly informative and may harbor genes for the quantitative traits. So far, the current statistical methods for QTL hotspot detection use either the individual-level data from the genetical genomics experiments or the summarized data from public QTL databases to proceed with the detection analysis. These detection methods attempt to address some of the concerns, including the correlation structure among traits, the magnitude of LOD scores within a hotspot and computational cost, that arise during the process of QTL hotspot detection. In this article, we describe a statistical framework that can handle both types of data as well as address all the concerns at a time for QTL hotspot detection. Our statistical framework directly operates on the QTL matrix and hence has a very cheap computation cost, and is deployed to take advantage of the QTL mapping results for assisting the detection analysis. Two special devices, trait grouping and top γ_*n,α*_ profile, are introduced into the framework. The trait grouping attempts to group the closely linked or pleiotropic traits together to take care of the true linkages and cope with the underestimation of hotspot thresholds due to non-genetic correlations (arising from ignoring the correlation structure among traits), so as to have the ability to obtain much stricter thresholds and dismiss spurious hotspots. The top γ_*n,α*_ profile is designed to outline the LOD-score pattern of a hotspot across the different hotspot architectures, so that it can serve to identify and characterize the types of QTL hotspots with varying sizes and LOD score distributions. Real examples, numerical analysis and simulation study are performed to validate our statistical framework, investigate the detection properties, and also compare with the current methods in QTL hotspot detection. The results demonstrate that the proposed statistical framework can effectively accommodate the correlation structure among traits, identify the types of hotspots and still keep the notable features of easy implementation and fast computation for practical QTL hotspot detection.

## INTRODUCTION

The quantitative trait loci (QTL) mapping experiments have been performed for traditional traits (such as yield and quality in rice, weight and body fat percentage in animals, diabetes and hypertensions in human) and molecular traits (such as gene expression or protein levels using the newly developed microarray technique) to explore the genetic mechanisms of these traits in various organisms and many areas of biological studies. When performing on traditional traits, a single experiment can produce abundant marker genotypes but usually consider only a few traits, say about 10~20 traits, in the population, since measuring traditional traits can be time-consuming and a costly process. On the contrary, when the experiment is conducted on molecular traits (called the genetical genomics experiment), with the aid of a high-throughput molecular biology techniques, it can not only produce abundant marker genotypes, but also generate thousands of molecular traits for the individuals at a time (Jansen and Nap 2001; Brem *et al*. 2002). To detect the QTL for these traits, many statistical methods have been proposed to analyze the QTL mapping data for the estimation of QTL parameters, including the QTL effects and positions, epistasis among QTL, heritabilities, etc. (Lander and Botstein 1989; Haley and Knott 1992; Jansen 1993; Zeng 1994; Kao *et al.*1999; Sen and Churchill 2001; Xu 2003; Broman *et al.* 2003; Kao 2006; Lee *et al.* 2014; Wei and Xu 2016; da Silva Pereira *et al*. 2020). In QTL mapping for either the traditional or molecular traits, it has been observed that QTL are highly abundant in some of the genomic regions, and that QTL responsible for correlated traits are frequently clustered closely together in some specific genetic regions as compared to other regions (Goffinet and Gerber 2000; Schadt *et al.* 2003; Chardon *et al.* 2004; West *et al.* 2007; Breitling *et al.* 2008; Wu *et al.* 2008; Wang *et al.* 2014; Basnet *et al.* 2015; Yang *et al.* 2019). These regions enriched in QTL are referred to as QTL hotspots, and, statistically, they harbor a significantly higher number of QTL than expected by random chance. It has been noted that the phenomenon of QTL hotspots may have several causes, such as: QTL with large and consistent effects can be identified in the similar regions under different conditions in various studies; QTL with high allelic polymorphisms have a greater chance of being detected in different crosses and environments; pleiotropic or closely linked QTL that control correlated traits are frequently co-localized in the same regions in different experiments (Falconer and Mackay 1996; Zhao *et al.* 2011; Vuong *et al.* 2015; Mengistu *et al.* 2016; Zhang *et al*. 2019). As the QTL hotspots can lead to identifying genes that affect the traits of interest, and further help to build networks among QTL hotspots, genes and traits, the QTL hotspot detection analysis at genome-wide level has been a key step towards deciphering the genetic architectures of quantitative traits in genes, genomes and genetics studies (Breitling *et al.* 2008; Fu *et al.* 2009; Neto *et al.* 2012; Wang *et al.* 2014; Yang *et al.* 2019).

Genome-wide QTL hotspot detection first needs to collect data with many QTL to proceed with the detection analysis. So far, both the genetical genomics experiments and public QTL databases can provide the data sets with many QTL for the hotspot analysis, but note that these two data sources have different structures. The genetical genomics experiment contains individual-level data (containing the original marker genotypes and many molecular traits) that allow to detect thousands of QTL in a single experiment. And public database (such as GRAMENE, Q-TARO, Rice TOGO browser, PeanutBase and MaizeGDB) curates thousands of summarized QTL data (containing the detected QTL, trait names and reference sources without any individual-level data) for various traditional traits from numerous independent QTL experiments. Using these two types of data, several statistical methods mainly based on permutation tests have been proposed to detect QTL hotspots. West *et al.* (2007), Wu *et al.* (2008), Li *et al.* (2010), Breitling *et al.* (2008) and Neto *et al.* (2012) developed statistical methods to detect QTL hotspots using the genetical genomics experiments. The methods of West *et al.*, Wu *et al.* and Li *et al.* (the *Q*-method) first perform QTL mapping at all genomic positions for all traits to construct a QTL matrix, whose column dimension has the same length as the genome size and row dimension corresponds to the number of traits. Then, they permuted the row elements of the QTL matrix separately by traits to compute the hotspot size thresholds for assessing the significance of QTL hotspots. As these methods do not account for the correlation structure among traits, the thresholds are severely underestimated, leading to the detection of too many spurious hotspots (Breitling *et al.* 2008; Neto *et al.* 2012;). To consider the correlation structure among traits, Breitling *et al.* (2008) permuted the individual-level data by shuffling the all traits relative to genotypes to generate numerous permuted data sets, and then performed QTL mapping on each of the permuted data sets to obtain the QTL matrices and determine the hotspot thresholds in hotspot detection. The way of permutation in Breitling *et al.* (the *N*-method) can preserve the correlation structure among traits and provide stricter thresholds to prevent spurious hotspots due to non-genetic correlation, but still it may neglect small and moderate sizes of hotspots with larger LOD scores (Neto *et al.* 2012). To further consider the LOD scores of QTL in hotspot detection, Neto *et al.* (2012) adopted the same permutation schemes and took the magnitude of LOD scores into account to compute a series of LOD thresholds for different hotspot sizes. The approach of Neto *et al.* (the *NL*-method) can effectively discover the small and moderate hotspots with strong LOD scores in QTL hotspot detection. Later, using the summarized QTL data in public databases, Yang, Wu and Kao (2019) proposed a statistical procedure to tackle the issue in genome-wide QTL hotspot detection. Briefly, the Yang *et al.* procedure first summarizes the data into a QTL matrix and converts it into an EQF (expected QTL frequency) matrix and further into a reduced EQF matrix by trait grouping. The trait grouping aims to account for the correlation structure among traits by grouping the (genetically) correlated traits together in the detection analysis. A permutation algorithm with trait grouping is then proposed to compute a series of EQF thresholds, from liberal to conservative, for assessing the significance of QTL hotspots. In this way, the Yang *et al.* procedure can cope with the underestimation of threshold due to ignoring the correlation structure among traits, and prevent detecting spurious hotspots. As well noted, the methods by permuting the individual-level data (the *N*- and *NL*-methods) involve repeated the QTL mapping analysis in each permutation and will suffer the problem of computational intractability, and may require parallel computations to complete the analyses (Neto *et al*. 2012). On the contrary, the methods by permuting the QTL (or EQF) matrix (the *Q*- and Yang *et al*. methods) need to perform QTL mapping analysis only once in the whole procedure, and therefore can offer the advantage of easy implementation and very cheap computation for QTL hotspot detection (Yang *et al.* 2019).

In this article, we introduce a general statistical framework that can handle both types of data as well as take care of all the above concerns, including the correlation structure among traits, the magnitude of LOD scores (a reflection of the QTL effects and their SD) in a hotspot and computational cost, for QTL hotspot detection. Our statistical framework operates on the QTL matrix or the EQF matrix and hence is very cheap in computation. By taking the advantages of using the individual-level data in the genetical genomics experiment, the estimates of QTL parameters and the LOD scores at every position for all traits can be obtained by QTL mapping and used to benefit the QTL hotspot analysis. Our statistical framework attempts to take the QTL mapping results into account to address the concerns and facilitate the hotspot detection. Two special devices, trait grouping and top γ_*n,α*_ profile, are deployed in the framework. In trait grouping, we show that, depending on the relative sizes and directions of QTL effects and the distances between QTL, the traits with tightly linked or pleiotropic QTL may have arbitrary values at their phenotypic or genetic correlations. Therefore, the estimated QTL positions, rather than the phenotypic or genetic correlations among traits, are used to directly make inference about the tightly linked and/or pleiotropic traits (the true linkages among QTL) for trait grouping, accounting for the correlation structure among traits. Then, the permutation algorithm of Yang *et al.* (2019) with trait grouping is deployed to compute a series of EQF thresholds, γ_*n,α*_’s, to assess the significance of QTL hotspots. For each hotspot, we profile the top γ_*n,α*_ thresholds and use the profile to outline the inside LOD-score pattern across the different LOD thresholds. The top γ_*n,α*_ profile can then serve to characterize the types of hotspots with varying sizes and LOD-score distributions, so as to have the ability to assess the small and moderate hotspots with strong LOD scores. In this way, our framework can overcome the underestimation of threshold arising from ignoring the correlation structure among traits, and also identify the different types of hotspots with very low computational cost during the process of hotspot detection. Numerical analysis, simulation study and real examples are conducted to explore the patterns of genetic correlations between closely linked and/or pleiotropic traits, investigate the properties of the proposed statistical framework, and assess the performances and compared with the current approaches. We demonstrate that the proposed statistical framework can deal with both types of data, effectively accommodate the correlation structure among traits, hotspot sizes, LOD-score distributions and still keep the notable features of easy implementation and fast computation for QTL hotspot detection.

## A FRAMEWORK FOR DETECTING QTL HOTSPOTS

Our statistical framework aims to operate on both the summarized data from public databases and individual-level data from the genetical genomics experiment for QTL hotspot detection. The key and basic idea of our framework is to perform permutation analysis on the QTL matrices or EQF matrices, rather than on the original individual-level data, and to well utilize the QTL mapping results in the process of QTL hotspot detection. Since the summarized data are an intermediate component in the detection process of using the original individual-level data, the framework for operating on the individual-level data to detect the QTL hotspots is first described, and that for summarized data will follow without additional treatment. In the following, how the QTL matrices are constructed from the different LOD thresholds in QTL mapping using the individual-level data is first described. Then we convert the QTL matrices into the EQF matrices by assuming that the QTL position is normally distributed over its own QTL interval. After that, we show that trait grouping on the base of the estimated QTL positions is more effective than that of the phenotypic or genetic correlations among traits in making inference about the true linkage among traits (in combining the closely linked or pleiotropic traits). By grouping together the traits affected by the tightly linked and/or pleiotropic QTL and treating them as a permutation unit, the permutation algorithm of Yang *et al*. (2019) is applied to the QTL matrices or EQF matrices and each computes a series of EQF thresholds, γ_*n,α*_’s, varying from strict to liberal, to assess the significance of the QTL hotspots. For every hotspot, we profile the top γ_*n,α*_’s thresholds across the different EQF matrices and use the top γ_*n,α*_ profile to identify the different types of QTL hotspots with varying sizes and LOD score distributions. Numerical analysis, simulation study and real examples are followed to validate our statistical framework, investigate the detection properties, and also compare with the current methods in QTL hotspot detection.

### LOD scores and QTL matrices

A variety of QTL mapping methods have been proposed to estimate the genetic architecture parameters of quantitative traits (see **INTRODUCTION**). The estimation is generally based on the likelihood principle. Using these methods, for each trait, a (partial) likelihood-ratio test (LRT) statistic or a LOD score (1 LOD≈4.6 LRT) is calculated to test for the existence of a QTL at every genomic position. The LOD scores at every position for all traits can be recorded into a LOD score matrix. Then, given a predetermined LOD threshold for the test, the LOD score matrix can be converted into a QTL matrix by assigning 1 to the detected QTL positions and 0 otherwise. To determine the appropriate LOD threshold, several analytical, computational and empirical approaches have been developed (Lander and Botstein 1989; Churchill and Doerge 1994; Piepho 2001; Chang *et al.* 2009; Guo 2011; Kao and Ho 2012). Lander and Botstein (1989) suggested using a LOD threshold of about 2.7 at 5% significance level for a single trait in a ≈1,000-cM tomato genome based upon simulated data. Kao *et al.* (1999) used a LOD threshold of about 2.61 at 5% level for each of the three traits in a ≈1,679-cM pine genome with 120 markers based on the Bonferroni adjustment. For the same pine data, a LOD threshold of about 2.70 at 5% level was obtained based on the Gaussian stochastic process (Kao and Ho 2012), and that of about 2.54 at 5% level was obtained by using the permutation test (Churchill and Doerge 1994) in our practice. We obtained a LOD threshold of about 3.06~3.49 at 5% level for the yeast data containing 6348-cM genome with 2956 markers (Brem and Kruglyak 2005) based on the Gaussian stochastic process (Guo 2011). The Gaussian process is found to be about 7700 times faster than the permutation test in obtaining the thresholds. Wang *et al*. (2014) applied the permutation test to the 100 e-traits that randomly selected from the 21,929 e-traits and obtained an LOD threshold of 4.76~5.10 at 5% level for the rice 1625-cM genome with 1619 markers. In general, obtaining thresholds using the analytical approaches, such as the Gaussian process, has a very cheap computational cost and may require the assumption of normal distribution, and that using the permutation test is robust to departures from normal assumption and needs to handle the problem of computational intractability. The LOD thresholds vary and increase with marker density, genome size and number of traits. Higher LOD thresholds yield less detected QTL of large sizes (high LOD scores). Therefore, in the context of hotspot detection analysis, for ease of computation, it is possible that the LOD thresholds can be determined based on empirical experience or by using the Gaussian process without bothering to use the permutation tests. Given a LOD threshold, the estimated QTL positions and their confidence intervals (constructed by using the asymptotic SD or LOD support intervals) can be obtained. For the summarized QTL data in the public databases, usually neither LOD scores nor the confidence intervals of QTL positions are reported, and only the flanking markers of the detected QTL are recorded. Here, we called the confidence interval of a QTL position or the marker interval containing a QTL a QTL interval. Then, we use a QTL matrix (an atypical matrix) with column dimension equivalent to the genome size and row dimension corresponding to the number of traits to summarize the QTL intervals for all traits as follows: For each trait, we mount the QTL intervals onto the elements of a row array as follows: Each QTL interval stands for an element of the length as its width at the corresponding position, and a value of one is given to the element. The remaining elements will be treated as zeros. Combining the arrays for all traits will form a QTL matrix, whose elements are either one or zero with unequal lengths (see Figure 1 in Yang *et al.* 2019 for graphical illustration). In this way, for a range of QTL mapping LOD thresholds from relaxed to conservative, say the LOD thresholds of 3 to 8 (by one increment), we can construct several (six) corresponding QTL matrices for operation. The natural choices for the relaxed and conservative LOD thresholds are the single-trait QTL mapping threshold controlling genome-wide error rate (GWER) for one trait and a multiple single-trait QTL mapping threshold controlling GWER across all traits, respectively, as suggested by Neto *et al.* (2012). The QTL matrices constructed with higher LOD thresholds will contain less QTL but of larger LOD scores, so will the hotspot size thresholds. Such a property can be applied to consider both hotspot size and LOD score distribution in QTL hotspot detection as described below.

**Figure 1:**
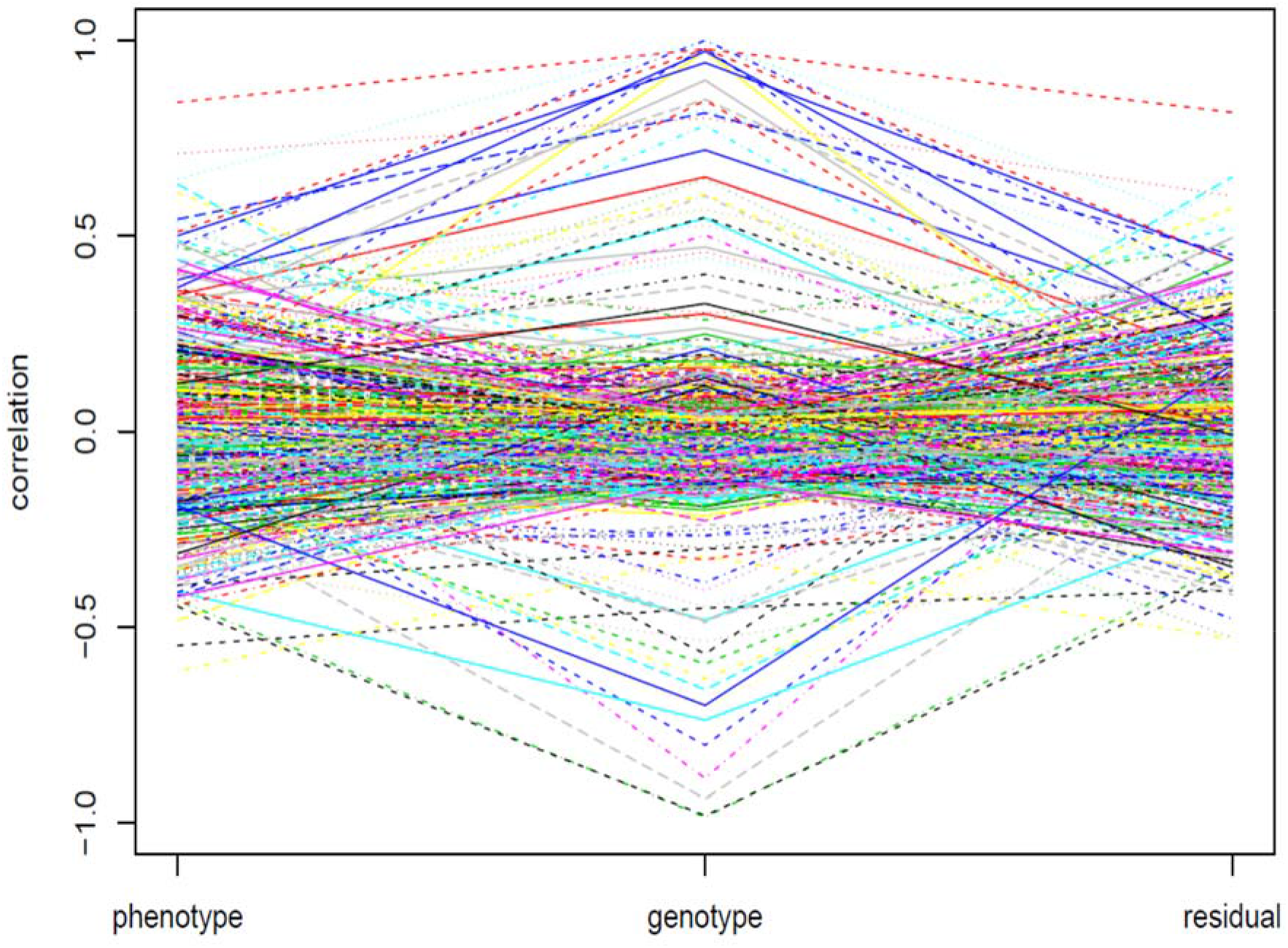
Phenotypic, genetic and residual correlations of the 500 randomly selected pairwise traits that controlled by the QTL with LOD scores larger than 3 in the yeast data set.

### Expected QTL frequency, EQF matrices and EQF architecture

We assume that *m* QTL matrices have been constructed from the LOD score matrix using the *m* different LOD thresholds (*L*_1_, *L*_2,_ ⋯, *L*_*m*_). We now take one QTL matrix as an example to show how to compute the expected QTL frequency (EQF) of a bin and to construct the corresponding EQF matrix, and the remaining EQF matrices from the other QTL matrices can be obtained in the same way. Consider that there are *T* traits each mapped for *N*_1_, *N*_2_,…, *N*_*T*_ QTL (intervals), respectively, where *N*= *N*_1_ + *N*_2_ + ⋯ + *N*_*T*_ is the total number of QTL. The genome is divided into *S* sequential equally spaced bins, each with the same size △ (say △=1 or 2 cM), for QTL hotspot analysis. For a bin (*x*, *x* + △) and a QTL interval (*a*, *b*), where *x*, *a* and *b* denote the genomic positions, they may have an overlap or no overlap. When there is an overlap, there is a probability that the QTL is localized in the bin, and the QTL will contribute a probability to the EQF value of this bin. Such a QTL will be referred to as a contributive QTL of a bin. We further assume that the QTL position is normally distributed over its own QTL interval to compute the contributed probability (EQF value) in a bin. Now let *f*_*ts*_ denote the EQF value of the *s*th bin between *x* and *x* +△ for the *t*th trait, where *t* = 1,2, ⋯, *T* and *s* = 1, 2, …, *S*. We have

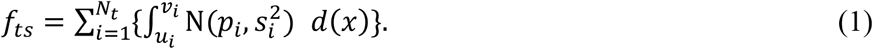

where *N*_*t*_ is the number of contributive QTL of the *t*th trait, (*u*_*i*_, *v*_*i*_) is the overlap region, *p*_*i*_ is the estimated QTL position (LOD peak) of the contributed QTL, and 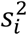 is the asymptotic variance. The asymptotic variance 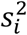 can be obtained in two way: (1) 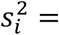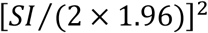, where *SI* is the 95% empirical support interval (Visscher *et. al.* 1996; Lynch and Walsh 1998); (2) 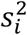 can be obtained by the general formulas of Kao and Zeng (1997). For each contributed QTL, a cumulative normal distribution probability ranging from *u*_*i*_ to *v*_*i*_ is added to the EQF of the bin. In general, the closer the overlap is to the LOD peak or the higher the LOD score of a QTL is, the greater the contributed probability is. Note that, for one QTL interval, our method using equation (1) assigns a fraction (the fractions of the within bins sum to one) and the *NL*-method assigns one to each of the within bins. For the QTL data from public databases, only the flanking markers are available for the QTL intervals, and the uniform distribution will replace the normal distribution for computing the EQF value (Yang *et al.* 2019). The EQF value can be calculated at each bin for each single trait to produce the EQF matrix as **F** = {*f*_*ts*_}_*T*×*S*_. The sum over the EQF values of all traits, *i.e.* 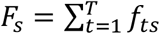, at every bin will produce the EQF architecture of the genome (see Figures 2, 3 or 4). A higher EQF value implies a greater expectation of localizing a QTL in the bin. A hotspot detection is claimed in the bin if its EQF value is higher than an EQF threshold that will be determined by permutation tests (see below).

**Figure 2.**
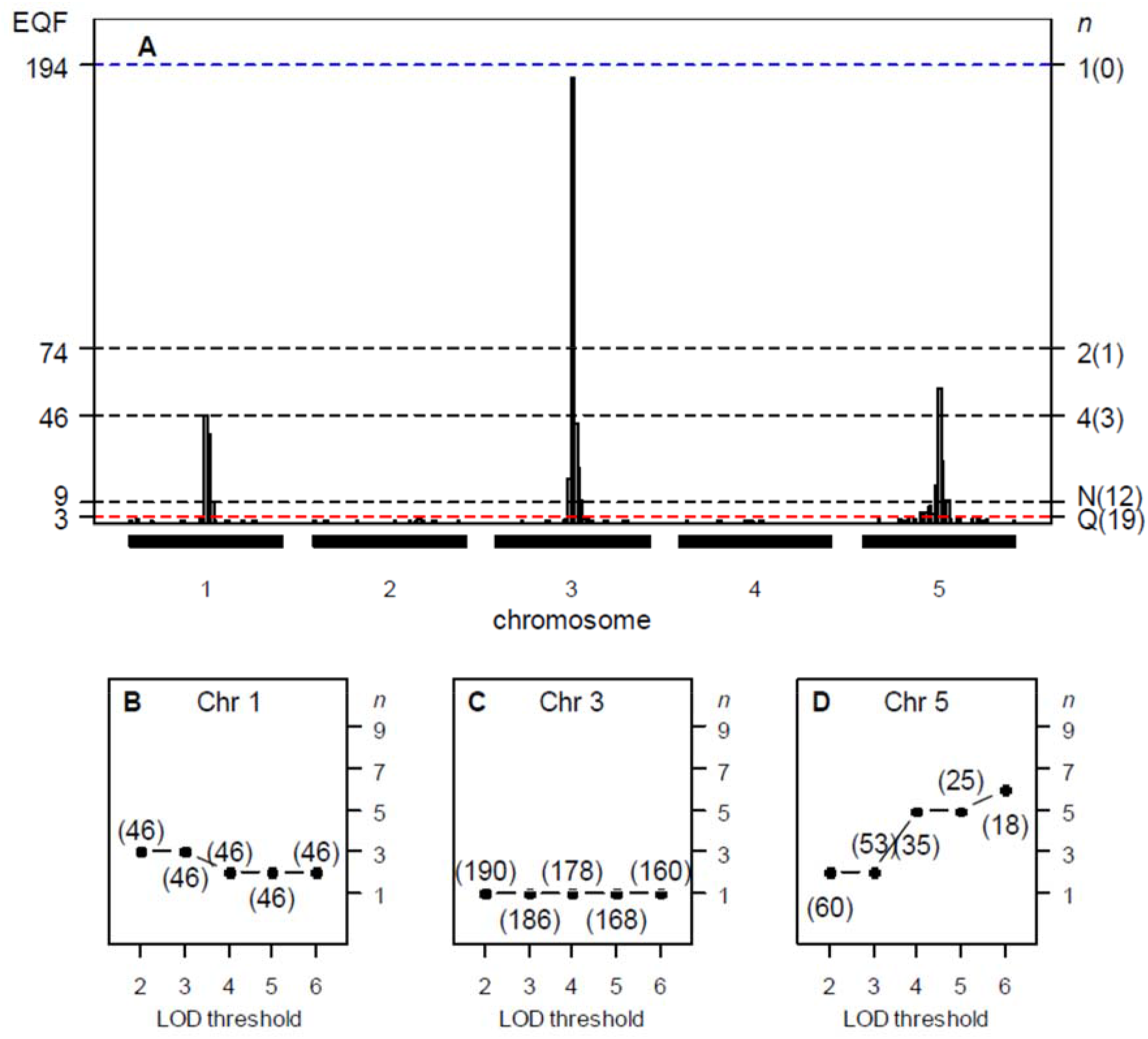
Panels (A-D) The hotspot architecture and the top γ_*n,α*_ profiles of the three simulated hotspots across the 2- to 6-LOD EQF architectures. Panels (A) Inferred hotspot architecture using a single-trait permutation LOD threshold of 2.47 corresponding to a GWER of 5% of falsely detecting at least one QTL somewhere in the genome. The hotspots on chromosomes 1, 3 and 5 have sizes 46, 188, and 57, respectively. The dotted line at count 9 corresponds to the hotspot size threshold at a GWER of 5% according to the *N*-method. The dash line at count 3 corresponds to the *Q-*method’s 5% significance threshold. The thresholds γ_1,0.05_, γ_2,0.05_ and γ_4,0.05_ obtained by the proposed procedure are 194, 74 and 46, respectively. Panel (B) The top γ_*n,α*_ profile for hotspot A shows a decreasing pattern with the value of *n* decreasing from 3 to 2 over the 2- to 6-LOD EQF architectures, indicating that hotspot A contains relatively more QTL with large LOD scores. (46,3): the size and top γ_*n*,0.05_ threshold for the hotspot is 46 and γ_3,0.05_, respectively. Panel (C) The top γ_*n,α*_ profile for hotspot B shows a flat pattern with *n*=1 for all the EQF architectures, indicating that hotspot B containing QTL with balanced LOD scores. Panel (D) The top γ_*n,α*_ profile for hotspot C shows an increasing pattern with the value of *n* increasing from 2 to 6 over the five EQF architectures, indicating that hotspot C contains relatively less QTL with large LOD scores. The number in the bracket is the number of detected hotspots. Results are based on 1000 permutations. Q: The *Q-*method; N: The *N*-method.

**Figure 3.**
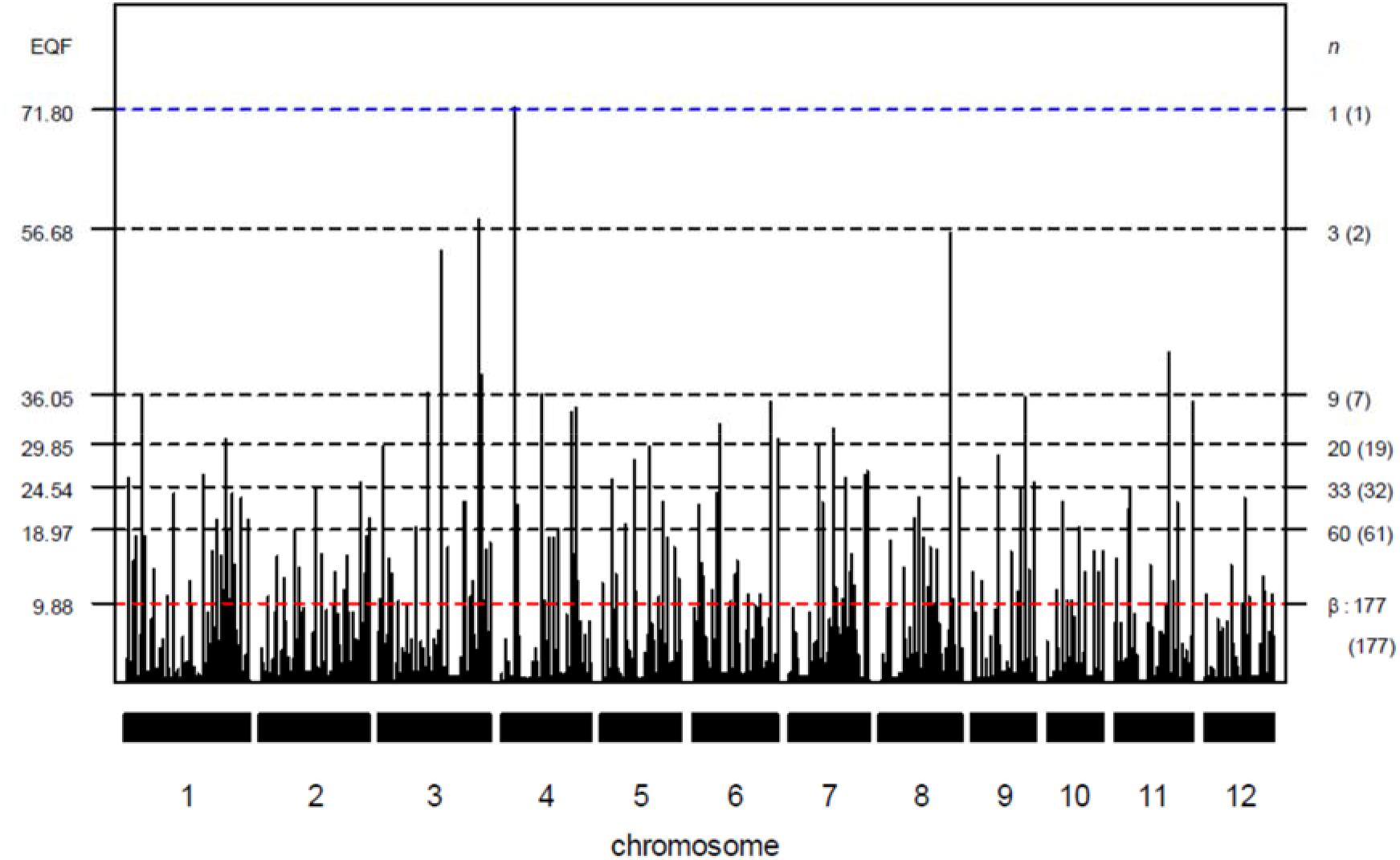
The EQF architectures along the 12 chromosomes and the hotspots detected under different EQF thresholds (γ_*n*,0.05_) associated with their qFreq(*n*) statistics at GWER of 5%. The thresholds γ_*n*,0.05_ are coordinately represented by the left and right axes. The left axis denotes the values of EQF, and the right axis denotes the values of *n*. The blue line corresponds to the EQF threshold γ_1,0.05_ = 71.80 for detecting at least one hotspot, and there is one (the number in the bracket) hotspot detected with γ_1,0.05_. Similarly, γ_3,0.05_ = 56.68 for detecting at least three hotspot, and there is two hotspot detected with γ_3,0.05_. The red line shows γ_177,0.05_ = 9.88 for detecting at least 177 hotspots, which approximately corresponds to *β* (the EQF threshold of the *Q*-method), and there are 177 significant hotspots with γ_177,0.05_.

**Figure 4.**
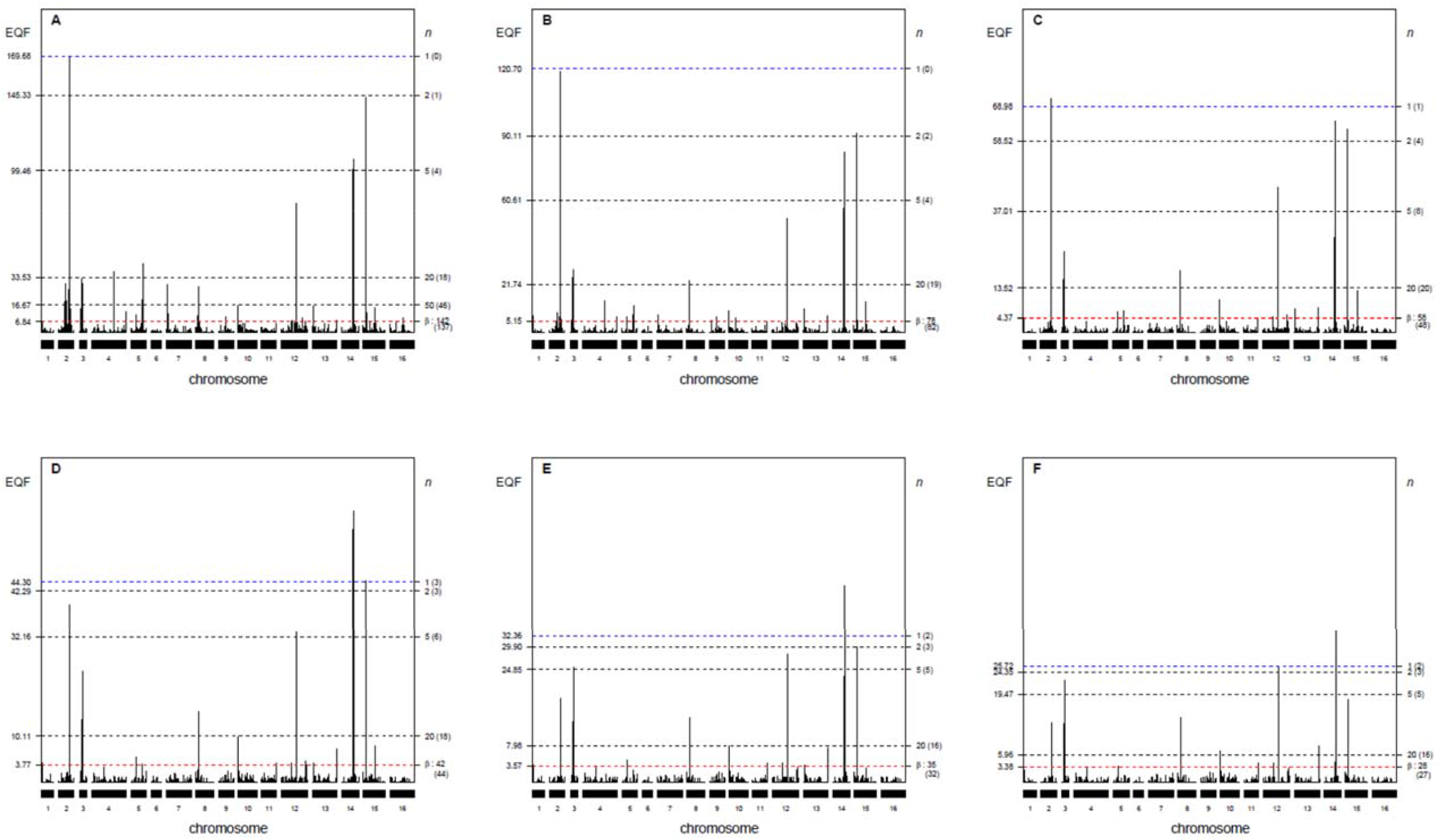
Panels (A-F) The EQF architectures along the 16 chromosomes and the hotspots detected under different EQF thresholds (γ_*n*,0.05_) at GWER of 5% in the 3− to 8-LOD EQF architectures with bin size of 2 cM. The left axis denotes the values of EQF, and the right axis denotes the values of *n*. The blue lines (first horizontal lines) correspond to the EQF thresholds γ_1,0.05_ = 169.68, 120.70, 68.98, 44.30, 32.36 and 25.72 for detecting at least one hotspot, and in practice there is 0, 0, 1, 3, 2, 2 hotspots detected with these γ_1,0.05_′s in the 3- to 8-LOD EQF architectures. The bottom horizontal (red) lines show the EQF thresholds of the *Q*-method, *β*’s, which approximately correspond to γ_142,0.05_, γ_78,0.05_, γ_58,0.05_, γ_42,0.05_, γ_35,0.05_ and γ_28,0.05_, respectively (see text). The number in the bracket is the number of detected hotspots.

### Trait grouping

Yang *et al*. (2019) suggested grouping the genetically correlated traits together to account for the correlation structure among traits and overcome the underestimation of threshold, preventing spurious hotspots. The primary purpose of grouping the genetically correlated traits together is to combine the traits controlled by the tightly linked and/or pleiotropic QTL together, and then treat them as a permutation unit in the permutation analysis to obtain stricter thresholds for QTL hotspot detection. Here, instead of using the phenotypic or genotypic correlations among traits, we use the estimated QTL positions directly to make inference about those tightly linked and/or pleiotropic traits for trait grouping. The reason is that using phenotypic or genetic correlations among traits is not effective and sufficient to group together the tightly linked and/or pleiotropic traits (the true linkage among QTL) as shown below. Let *P*, *G* and *E* denote the phenotypic value, genotypic value and residual of a quantitative trait, respectively, then we have *P* = *G* + *E* as usual. For a pair of traits, *P*_1_ and *P*_2_, we can derive

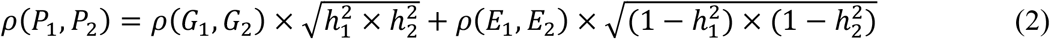

(see also Falconer and Mackay 1996), where *ρ*(*P*_1_, *P*_2_), *ρ*(*G*_1_, *G*_2_) and *ρ*(*E*_1_, *E*_2_) are the phenotypic, genetic and residual correlations between the two traits, respectively, and 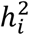 is the heritability. Equation (2) tells that phenotypic correlation between two traits is an outcome of interplays among genetic correlation, residual correlation and heritability. Some real examples of the three correlations between traits include: Zeng, Kao and Basten (1999) analyzed the cone number and branch quality in pine and found that phenotypic correlation is very small (0.013), the genetic correlation is estimated to be significantly negative (−0.196), and the residual correlation is estimated to be significantly positive (0.189). The heritabilities are 0.560 and 0.363, respectively. We analyzed the 5740 molecular traits in the yeast data (Brem and Kruglyak 2005), and found that there are various types of relationships between the three correlations. Figure 1 presents the phenotypic, genetic and residual correlations of 500 randomly selected pairs of traits in the yeast data set after performing the QTL mapping analysis. It shows that, for a pair of traits, high (low) phenotypic correlation does not imply high (low) genetic correlation, and vice versa, as the heritability and residual correlation also play roles in affecting the strength of phenotypic correlation. The phenotypic and genetic correlations seem to have an arbitrary relationship, making accurate prediction of each other difficult.

We further show that a pair of traits controlling by tightly linked QTL or pleiotropic QTL may have arbitrary values at their genetic correlation depending on the effects and linkage parameters using a backcross population. First, we consider the case of two monogenic traits. Assume that the first trait is affected by a QTL, Q_*i*_, and the second trait is affected by another QTL, Q_*j*_. We have *ρ*(*G*_1_, *G*_2_) = ±(1 − 2*r*_*ij*_), where *r*_*ij*_ is the recombination fraction between Q_*i*_ and Q_*j*_, despite of their effect sizes. The value of *ρ*(*G*_1_, *G*_2_) is positive (negative) if they have the same (opposite) direction of effects. If the two traits are pleiotropic (*r*_*ij*_ = 0), we have *ρ*(*G*_1_, *G*_2_) = ±1. Further, we consider that the two traits both have a digenic inheritance. Assume that the first trait is controlled by Q_*i*_ and Q_*k*_, and the second trait is controlled by Q_*j*_ and Q_*l*_. We have the model *G*_1_ = *μ*_1_ + *a*_1_*x*_*i*_ + *a*_2_*x*_*k*_ to model *G*_1_, where *x*_*i*_ and *x*_*k*_ are coded variables for Q_*i*_ and Q_*k*_, and *a*_1_ and *a*_2_ are their effects. Similarly, we have the model *G*_2_ = *μ*_2_ + *b*_1_*x*_*j*_+*b*_2_*x*_*l*_ to model *G*_2_, where *x*_*j*_ and *x*_*l*_ are coded variables for Q_*j*_ and Q_*l*_, and *b*_1_ and *b*_2_ are the effects. For the case of Q_*i*_-Q_*j*_---Q_*k*_-Q_*l*_ order, the genetic correlation between the two digenic traits is

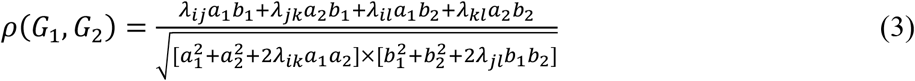

where *λ*_*ij*_ = 1 − 2*r*_*ij*_ is the linkage parameter between Q_*i*_ and Q_*j*_, showing that the genetic correlation is a function of QTL effects and their linkage parameters. To simplify the analysis of equation (3), we consider the case that the QTL are pleiotropic and unlinked (Q_*i*_=Q_*j*_, Q_*k*_=Q_*l*_, *λ*_*ij*_ = *λ*_*kl*_ = 1, *λ*_*ik*_ = *λ*_*il*_ = *λ*_*jk*_ = *λ*_*jl*_ = 0.5) so that equation (3) reduces to

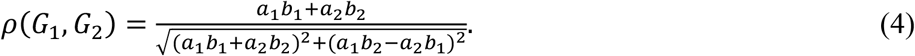

Bases on equation (4), some examples of arbitrary genetic correlations between the two digenic pleiotropic traits are: If (*a*_1_ = 5, *a*_2_ = 2, *b*_1_ = 2, *b*_2_ = −5), we have *a*_1_*b*_1_ + *a*_2_*b*_2_ = 0 and *ρ*(*G*_1_, *G*_2_) = 0. If (*a*_1_ = 2, *a*_2_ = 2, *b*_1_ = 2, *b*_2_ = 2), we have *a*_1_*b*_2_ − *a*_2_*b*_1_ = 0 and *ρ*(*G*_1_, *G*_2_) = 1. If (*a*_1_ = 2, *a*_2_ = 2, *b*_1_ = 2, *b*_2_ = 2), we have *a*_1_*b*_2_ − *a*_2_*b*_1_ = 0 and *ρ*(*G*_1_, *G*_2_) = −1. If (*a*_1_ = 5, *a*_2_ = 2, *b*_1_ = 2, *b*_2_ = 5), we obtain *ρ*(*G*_1_, *G*_2_) = 0.690. If (*a*_1_ = 5, *a*_2_ = 2, *b*_1_ = −2, *b*_2_ = −5), we obtain *ρ*(*G*_1_, *G*_2_) = −0.690. Therefore, depending on the relative sizes and directions of effects, their genetic correlations can be −1, −0.690, 0, 0.690 and 1, showing that the digenic pleiotropic traits may have very strong to very weak genetic correlations as well as no genetic correlation.

Table 1 presents the numerical analysis of equation (3) on the genetic correlation *ρ*(*G*_1_, *G*_2_) between the two digenic traits controlled by one or two pleiotropic QTL, respectively, under different settings of effect sizes and linkage parameters. For the case of two pleiotropic QTL, it shows that *ρ*(*G*_1_, *G*_2_) can be any value between 1 and +1 for different effect sizes, despite of being linked or unlinked. For example, if they are unlinked, the values of *ρ*(*G*_1_, *G*_2_) are 1, 0.690, 0, −0.690, 0, −0.690 and −1, respectively, for the seven effect size settings. The values are 1, 0.930, 0.547, −0.036, −0.547, −0.930 and −1, respectively, for the case of 20-cM apart. For the setting of (*a*_1_ = −2, *a*_2_ = 5, *b*_1_ = 5, *b*_2_ = 2), the values are 0.296, −0.036, −0.431 and −0.690, respectively, if the two pleiotropic QTL are 10-, 20-, 50-cM apart and unlinked, showing that a closer linkage between the pleiotropic QTL does not necessarily imply a higher genetic correlation. Similar pattern can be observed for the case of the digenic traits with one pleiotropic QTL. In general, the above analytical and numerical analyses indicate that, depending on the effect sizes and linkage parameters, pleiotropic traits may show no genetic correlation, or very weak to very strong genetic correlations. The relationship between pleiotropic traits and their genetic correlations is uncertain. As is well known, genetic correlations between traits are caused by the closely linked and pleiotropic QTL (Falconer and Mackay 1996). Nevertheless, here we present that multigenic traits controlled by tightly linked and pleiotropic QTL do not necessarily show significant genetic correlations with each other, because linkages between different QTL may individually make different levels of (positive or negative) contribution and may together combine to produce a low genetic correlation.

**Table 1.** The genetic correlations between two digenic traits controlled by one or two pleiotropic QTL under different settings of relative sizes and directions of QTL effects and the linkage parameters.

| $\begin{array}{c} \vert \quad d_{ik} \quad \vert \\ Q_i \quad (Q_j) \quad Q_k \quad (Q_l) \end{array}$ | | $(a_1, a_2, b_1, b_2)$ | | | | | | |
| --- | --- | --- | --- | --- | --- | --- | --- | --- |
| $d_{ik}$ | $d_{kl}$ | (2,2,5,5) | (2,5,5,2) | (-2,5,5,2) | (-2,5,5,-2) | (2,-5,5,2) | (2,5,-5,2) | (2,-2,-5,5) |
| 10 | - | 1.000 | 0.964 | 0.718 | 0.296 | -0.718 | -0.964 | -1.000 |
| 20 | - | 1.000 | 0.930 | 0.547 | -0.036 | -0.547 | -0.930 | -1.000 |
| 50 | - | 1.000 | 0.843 | 0.275 | -0.431 | -0.275 | -0.843 | -1.000 |
| ∞ | - | 1.000 | 0.690 | 0.000 | -0.690 | 0.000 | -0.690 | -1.000 |

| $\begin{array}{c} \vert \quad d_{ik} \quad \vert \quad d_{kl} \quad \vert \\ Q_i \quad (Q_j) \quad Q_k \quad (Q_l) \end{array}$ | | $(a_1, a_2, b_1, b_2)$ | | | | | | |
| --- | --- | --- | --- | --- | --- | --- | --- | --- |
| $d_{ik}$ | $d_{kl}$ | (2,2,5,5) | (2,5,5,2) | (-2,5,5,2) | (-2,5,5,-2) | (2,-5,5,2) | (2,5,-5,2) | (2,-2,-5,5) |
| 10 | 10 | 0.949 | 0.942 | 0.690 | 0.354 | -0.690 | -0.942 | -0.949 |
| 20 | 20 | 0.897 | 0.878 | 0.499 | 0.105 | -0.499 | -0.878 | -0.897 |
| 50 | 50 | 0.751 | 0.690 | 0.089 | -0.165 | -0.089 | -0.690 | -0.751 |
| ∞ | ∞ | 0.500 | 0.345 | -0.345 | -0.345 | 0.345 | -0.345 | -0.500 |

The first trait is controlled by $Q_i$ and $Q_k$ with effects $a_1$ and $a_2$ , the second trait is controlled by $Q_j$ and $Q_l$ with effects $b_1$ and $b_2$ . $d_{ik}$ and $d_{kl}$ denote the genetic distances between $Q_i$ and $Q_k$ and between $Q_k$ and $Q_l$ , respectively.

The above analyses of equations (2), (3) and (4) tell us that (a) phenotypic correlation between traits may have no relationship with their genetic correlation, (b) the genetic correlation between monogenic traits controlled by a pleiotropic QTL is either −1 or +1, (c) depending on their effect sizes and linkage parameters, the genetic correlations between polygenic traits with one or more pleiotropic QTL may show very weak to very strong genetic correlations, ranging between −1 to +1, including no genetic correlations, and (d) trait grouping based on the genetic correlations can only combine the closely linked or pleiotropic traits with high genetic correlations, but will fail to combine those with weak (or no) genetic correlations. Based on these findings, instead of using the genetic correlation, we suggest directly using the estimated QTL positions (significant LOD peaks) to make inference about the true linkage among traits for trait grouping. For example, we can define those QTL being localized in the same bins (*e.g.*, bin size of 0.5, 1 or 2 cM) as tightly linked and/or pleiotropic QTL, and group their traits together as trait groups (hereinafter referred to as “the empirical trait grouping”). The traits in each trait group will be treated as a trunk or unit of permutation in the analysis to cope with the correlation structure among traits and obtain stricter thresholds for assessing the significance of QTL hotspots.

### Permutation algorithm for computing the EQF thresholds γ_*n,α*_’s

We devise two permutation schemes, the QTL-interval permutation and the EQF-bin permutation, to compute the EQF thresholds for assessing the significance of QTL hotspots. The QTL-interval permutation operates on the QTL matrix, and then considers the genome to be circular (Cabrera *et al.* 2012) and randomly swaps the QTL intervals in the circular genome. On the contrary, the EQF-bin permutation works on the EQF matrix, and then breaks a QTL interval into several EQF bins and randomly shifts the EQF bins along the genome. When trait grouping is considered to cope with the correlation structure among traits, the QTL intervals or the EQF bins in each trait group will be permuted together, and the stricter EQF thresholds can be obtained to assess the hotspot significance. Below the EQF-bin permutation algorithm is first outlined, and the QTL-interval permutation algorithm will follow without much difficulty.

Using the *m* different LOD thresholds, we have converted the LOD score matrix into the *m* QTL matrices and then *m* EQF matrices, respectively. For each of the *m* EQF matrix, **F**, we first obtain the EQF sum over all traits for the *S* bins and order them from highest to lowest, *F*_(1)_, *F*_(2)_, …, *F*_(*s*)_. Then, we define qFreq(*n*) as the *n*th EQF sum of the *S* ordered observed EQF sums, and use qFreq(*n*) as a test statistic for at least *n* spurious hotspots under the null hypothesis that the QTL are randomly distributed in the genome. The steps of the EQF-bin permutation algorithm with trait grouping to compute the threshold that can control GWER of qFreq(*n*) at a fixed α level is described below:

1. The traits with tightly linked or pleiotropic QTL are grouped together. After grouping, there are, say *R*, trait groups containing *g*_1_, *g*_2_, *g*_3_, ⋯, *g*_R_ traits, respectively (∑ *g*_*i*_ = *T*). Each trait group will be regarded as a permutation unit, *i.e.* the traits in the same group will be permuted together, in permutation analysis.
2. Generate a new permuted matrix **F**\* by performing permutation in each trait group as follows: For the *i*th trait group (*i* = 1, 2, ⋯, *R*), the *g*_i_ EQF values, *i.e.* the *g*_i_ row elements, at each bin are together randomly allocated across the *S* genomic locations.
3. Compute the total EQF sums over all rows for the *S* locations, *i.e.* 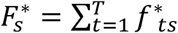 for *s* = 1, 2, …, *S*, for the **F**\* matrix, and order them 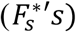 from highest to lowest as 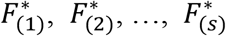.
4. For a fixed hotspot number *n*, obtain and store 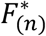 corresponding to the *n*th EQF sum of the *S* ordered EQF sums for **F**\*.
5. Repeat steps 1–4 *B* times so that there are *B* new permuted matrices (namely, **F**^1^, **F**^2^,…, **F**^*B*^) for obtaining the associated 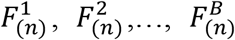. The *B* -permutation samples of 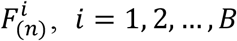, is an estimate of the null distribution of the test statistic qFreq(*n*) for at least *n* spurious hotspots anywhere in the genome under the null.
6. The upper (1-α)-quantile of the *B*-permutation samples generated in step 5 is the EQF threshold, denoted by γ_*n,α*_, for qFreq(*n*) for assessing at least *n* spurious hotspots.

The EQF-bin permutation with trait grouping can be also performed by first obtaining the reduced EQF matrix (with a dimension *R* × *S*, where 2 ≤ *R* ≤ *T*), which is constructed by pooling together the EQF values in each of the trait groups, *i.e.* by combining the row arrays of the same groups into a single row array, and then performing permutation analysis on the reduced EQF matrix to obtain the γ_*n,α*_ threshold (Yang *et al.* 2019). Besides, the above permutation algorithm can be also directly applied to the QTL-interval permutation as follows: First consider the genome to be circular and then permute together the QTL intervals in each trait group to obtain the permuted QTL matrices. For each of the permuted QTL matrices, the EQF sums of all bins (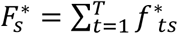 for *s* = 1, 2, …, *S*.) are computed using equation (1) and ordered from highest to lowest as 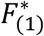,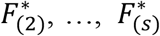 to obtain the γ_*n,α*_ threshold (step 3 to step 6). In this way, the proposed algorithm can deploy both the EQF-bin permutation and the QTL-interval permutation to compute a series of thresholds, γ_*n,α*_’s, for qFreq(*n*)’s to assess the significance of QTL hotspots. For *n* = 1, 2,⋯, *k*, where *k* is determined by *β* = γ_*k,a*_ (*β* is the threshold value obtained without trait grouping, *i.e.* by using the Q-method), a series of γ_*n,α*_’s ranging from the most conservative (*n* = 1) to the most liberal (*n* = *k*) can be obtained and used for assessing the significance of different numbers of QTL hotspots. By adopting γ_*n,α*_, it can control GWER of qFreq(*n*) at level α of detecting at least *n* spurious hotspots under the null, as detecting more than *n* hotspots is less likely than detecting *n* hotspots given the threshold γ_*n,α*_. For example, γ_1,*α*_ for detecting at least one hotspot is stricter than γ_2,*α*_ for detecting at least two hotspots, and a hotspot significant under γ_1,*α*_ must be also significant under γ_2,*α*_, and so on. As the permutation algorithm is performed on the QTL matrix or EQF matrix and only one QTL mapping analysis is required in the whole procedure, our framework has a very cheap computational cost as compared to the *N*-method and *NL*-method that need to perform many repeated QTL mapping analyses, and hence is suitable for practical use.

### The top γ_*n,α*_ profile

The permutation algorithm allows to compute a series of EQF thresholds, γ_*n,α*_’s, for each of the *m* EQF matrices. Now, we denote **F**(*L*_*i*_) as the EQF matrix constructed using the LOD threshold *L*_*i*_, and γ_*n,α*_(*L*_*i*_)’s as the corresponding series of thresholds for F(*L*_*i*_). We define the top γ_*n,α*_(*L*_*i*_) threshold for a bin *S* as

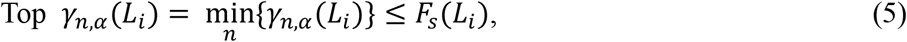

where *F*_*s*_(*L*_*i*_) is the EQF sum of the bin *S* over all traits in the **F**(*L*_*i*_) matrix. That is the top γ_*n,α*_(*L*_*i*_) threshold of the bin *S* is the largest EQF threshold (with the smallest *n*) for the bin to be significant as a QTL hotspot in the **F**(*L*_*i*_) matrix. For a hotspot, the smaller the value of *n* in its top γ_*n,α*_ threshold is, the relatively more significant as a hotspot it is. Therefore, in a specific EQF architecture, the top γ_*n,α*_ threshold of a hotspot can be used to characterize its significance status relative to the other hotspots. When considering across all the EQF architectures, there are *m* top γ_*n,α*_ thresholds for a hotspot. The pattern of (the *n* values in) the *m* top γ_*n,α*_ thresholds can outline how the relative significance status of the hotspot changes over the different EQF architectures. For example, suppose that, in the **F**(3) matrix, the bin *S* has an EQF value 43.89, which is significant under γ_10,0.05_ =43.12 (will be significant under γ_11,0.05_, γ_12,0.05_,⋯ also) but not significant under γ_,0.05_ =44.70. Then, the top γ_*n,α*_ threshold for the bin *S* is γ_10,0.05_(3) = 43.12 in **F**(3). Also, in the **F**(5) matrix, the bin *S* has a (smaller) EQF value, say 25.92, and the top γ_*n,α*_ threshold is γ_3,0.05_(5) = 25.1. The top γ_*n,α*_ thresholds for the bin *S* are γ_10,a_(3) and γ_3,*α*_(5), respectively, so the value of *n* changes from 10 in the **F**(3) matrix to 3 in the **F**(5) matrix, meaning that the bin *S* is significant as a hotspot under the γ_10,a_ threshold of detecting at least 10 hotspots in **F**(3) and under the γ_3,*α*_ threshold of detecting at least 3 hotspots in **F**(5). The bin *S* has a higher hotspot significance status in the 5-LOD EQF matrix than in the 3-LOD EQF matrix (*n* = 3 vs. *n* = 10), which implies that the QTL in the bin *S* has a relatively considerable amount of QTL with high LOD scores (LOD scores > 5) as compare to other bins. Therefore, by investigating how the value of *n* changes among the top γ_*n,α*_(*L*_*i*_) thresholds of a hotspot, we can understand the LOD score distribution of the hotspot relative to those of other hotspots across the different EQF architectures. In this way, we can compute and profile the top γ_*n,α*_ thresholds for a QTL hotspot in the different F(*L*_*i*_) matrices (say *L*_*i*_ = 3, 4, 5, 6, 7, 8, depending on the number of trait and genome size), and further use the top γ_*n,α*_ profile to outline the LOD score distribution of a hotspot. If the top γ_*n,α*_ profile of a hotspot shows a decreasing (increasing) pattern of *n* with *L*_*i*_ increasing, we may conclude that the hotspot contains relatively more (fewer) QTL with high LOD scores, as compare to other hotspots (see Figures 2 and 6). If the profile has a nearly flat pattern, we may infer that the QTL within the hotspot are well-balanced in the magnitude of the LOD scores. Based on the above interpretation, the pattern of the top γ_*n,α*_ profile of a QTL hotspot can readily identify and characterize the types of hotspots with varying sizes and LOD score distributions.

## SIMULATION STUDY AND REAL EXAMPLES

In this section, simulation study and real example analysis are conducted to illustrate the proposed statistical framework, investigate the related properties, and evaluate the performance as well as compare with the current methods in QTL hotspot detection. In simulation study, we investigate the performance of the proposed statistical framework and compare with the *Q*-method, the *N*-method and the *NL*-method in detecting QTL hotspots. In real example analysis, we first apply the statistical framework to analyze the summarized QTL data collected in GRAMENE rice database (Yang *et al.* 2019), and then to analyze the individual-level yeast data set in Brem and Kruglyak (2005) and compare the results with those in Yang *et al.* (2019) and Neto *et al.* (2012) in QTL hotspot detection, respectively. We also investigate and validate the patterns of genetic correlations among the closely linked and pleiotropic traits in the yeast data set to confirm the theoretical analysis in ***trait grouping***.

### Simulation study

Yang *et al.* (2019) have performed the simulation study to show that the permutation procedure with trait grouping can control GWERs at the target levels for the QTL data with correlation structure, and has the ability to produce quality results by offering a sliding scale of thresholds for QTL hotspot detection. In the Yang *et al.* simulation study, a perfect trait grouping, in which all the simulated pleiotropic traits are correctly grouped together, is considered for grouping traits. Here, using the same simulation settings, we now further show the effectiveness of the empirical trait grouping, in which the traits with QTL being localized in the same bins are grouped together, and investigate the ability of the top γ_*n,α*_ profile in characterizing and identifying the different types of QTL hotspots with varying sizes and LOD score distributions in the detection analysis. Likewise, we simulate a small-scale genetical genomics data set that contains 100 backcross progeny with 5 chromosomes of length 100 cM and 600 molecular traits. Each chromosome contains 50 equally spaced markers, and the bin size of 2 cM (Similar to that in Yang *et al.* 2019) is used in the analysis. We assume that all the 600 traits are monogenic. Each trait is controlled by one of the three unlinked genes, causing three hotspots A, B and C, respectively: (1) A small hotspot A is caused by a gene at 50 cM of the 1^st^ chromosome and is composed of 100 pleiotropic traits with heritabilities 0.3~0.45 showing high LOD scores in QTL mapping; (2) A big hotspot B is caused by a gene located at 50 cM of the 3^rd^ chromosome and contains 300 pleiotropic traits. Among the 300 pleiotropic traits, half have heritabilities 0.1~0.45 showing moderate to high LOD scores, and half have heritaibilities 0.3~0.45 showing high LOD scores; (3) A big hotspot C contains 200 traits and is caused by a gene at 50 cM of the 5^th^ chromosome. The heritabilities of the 200 traits are 0.1~0.2 showing small LOD scores. Figure S1, A-D, shows the LOD distributions of the traits in the three hotspots and the distribution of the pairwise correlation among traits for the simulated data set. The pairwise correlations among traits vary from −0.42 to 0.67 with mean 0.114 (Figure S1D). For the purpose of comparison, the bin containing the estimated QTL position will be given to 1 (and 0 otherwise) to construct the QTL matrix for the operation of the *Q*-method and our approach. The EQF-bin permutation and empirical trait grouping are adopted in the analysis. We also adopt 1.5-LOD 95% support intervals to decrease the spread of the hotspots for the *N*-method and *NL*-method (Neto *et al.* 2012).

The results of the *Q*-method, *N*-method, the *NL*-method and the proposed procedure (with empirical trait grouping) are summarized in Figure S2, A-F (see supplementary material), which are similar to those in Figure 6 of the Yang *et al*. paper using perfect trait grouping. Figure S2A and Figure 2 presents the hotspot architecture constructed using a single-trait LOD threshold of 2.47 (obtained by permutation test) and the 5% significance hotspot size thresholds obtained by the *Q*-method, *N*-method, and the proposed framework. The hotspots on the 1^st^, 3^rd^ and 5^th^ chromosomes have sizes 46, 188, and 57, respectively. The hotspot size thresholds obtained by the *Q*-method, *N*-method are 3 and 9, which correspond to γ_21,0.05_ and γ_13,0.05_ by our approach. Figures S2, B-F, present the hotspot architectures inferred using the *NL*-method LOD thresholds of 5.19, 3.77, 1.58, 1.36, and 1.13 that aim to control GWER of 5% for spurious hotspots of sizes 1, 3, 43, 60, and 90, respectively. It shows that, in addition to detecting the three true hotspots, the *Q*-method and *N*-method also detect several (19 and 12) spurious hotspots near the true hotspots due to lower thresholds, and the proposed procedure can produce less spurious hotspots due to higher thresholds (see Figure 2A). For example, using γ_2,0.05_ = 74 (the threshold for detecting at least two hotspots) as a threshold, only the hotspot on the 3^rd^ chromosome is detected (Figure 2A). Using γ_4,0.05_ = 46 (the threshold for detecting at least four hotspots) as a threshold, all the three hotspots can be detected (Figure 2A). It shows that the empirical trait grouping in our framework is effective to cope with the correlation structure among traits and can obtain stricter thresholds, preventing spurious hotspots in QTL hotspot detection. Figure 2, B-D, presents the top γ_*n,α*_ profiles of the three hotspots, A, B and C, in the 2- to 6-LOD EQF architectures. In Figure 2B, the top γ_*n,α*_ profile of hotspot A has a decreasing pattern with the value of *n* decreasing from 3 to 2 over the 2- to 6-LOD EQF architectures, indicating that hotspot A contains relatively more QTL with large LOD scores. In Figure 2C, the top γ_*n,α*_ profile for hotspot B has a flat pattern with *n*=1 for all the EQF architectures, indicating that hotspot B is a major hotspot containing QTL with balanced LOD scores. In Figure 2D, the top γ_*n,α*_ profile of hotspot C shows an increasing pattern with the value of *n* increasing from 2 to 6 over the five EQF architectures, indicating that hotspot C contains relatively less QTL with large LOD scores. Therefore, the patterns of the top γ_*n,α*_ profiles can outline the LOD-score distributions of the hotspots, A, B and C, well. To sum up, the simulation study shows that the proposed statistical procedure with trait grouping and the top γ_*n,α*_ profile has the ability to produce quality results by offering a sliding scale of thresholds from high to low for QTL hotspot detection, and is applicable to distinguish the different types of hotspots in the hotspot analysis.

### Real examples

#### The Gramene rice database

The GRAMENE database is a web-accessible and common reference database for crop research. For rice, it collects 8216 QTL (*N* = 8216) responsible for 236 different traits (*T* =236) from 230 published studies (experiments). The total length of the rice 12 chromosomes is ~1536.9 cM. There are 1914 common markers on the consensus map with an average marker density of one marker every 0.81 cM. The QTL density is ~5.35 QTL per cM. Among the 8216 QTL collected in the GRAMENE database, 309 (3.76%) QTL are localized at markers, 3791 (46.14%) QTL are localized in the marker intervals with sizes between 0 and 0.5 cM, 74 (0.90%) QTL are localized in the 0.5-1 cM intervals, 200 (2.43%) QTL are localized in the 1-2 cM intervals, 509 (6.20%) QTL are in the 2-5 cM intervals, 6.94 (5.57%) QTL are in the 5-10 cM intervals, and 1023 (12.45%) QTL are in the 10-20 cM intervals. The medium, mean and SD of the interval sizes are 0.56, 9.82 and 16.82 cM, respectively. It implies that if bins are identified as hotspots, the major contribution to their EQF is from the ≦1 cM QTL intervals, and that the large QTL intervals only contribute a small portion to the EQF of the hotspots. The flanking marker pairs of the 8216 QTL (8216 QTL intervals) are recorded and used for detecting QTL hotspots. By using equation (1) with uniform distribution and using bin size of 0.5 cM (△=0.5 cM), the EQF architectures (Figure 3) and the EQF matrix (with a dimension 236×3070) can be obtained. The EQF-bin permutation with empirical trait grouping is performed to obtain the EQF thresholds, γ_*n*,0.05_’s. When adopting the empirical trait grouping, only those QTL intervals ≦ 0.5 cM are considered, and there are a total of 71 trait groups in the analysis.

Figure 3 presents the EQF architectures of the 12 chromosomes and the hotspots detected under different EQF thresholds. In Figure 3, the 1^st^ highest peak (on the 4th chromosome) has an EQF value 71.97, and the threshold γ_1,0.05_ for detecting at least one QTL hotspot is 71.80. Under γ_1,0.05_ =71.80, only the 1^st^ highest peak was significant as a hotspot. Under γ_3,0.05_ =56.68 for detecting at least three hotspots, there are two hotspots detected (on the 3^rd^ and 4^th^ chromosomes). The highest peak of the 1^st^ chromosome has an EQF value 35.89 and was significant under γ_11,0.05_ =35.74, but not significant under γ_10,0.05_ = 35.90, and the top γ_*n,α*_ threshold associated with this peak is γ_11,0.05_. The EQF value of the 6^th^ highest peak (on the 3rd chromosome) is 38.52, and the top γ_*n,α*_ threshold is γ_7,0.05_ = 37.70. Under γ_7,0.05_ = 37.70, there are 6 significant hotspots in practice. Under γ_100,0.05_, there are 102 significant hotspots. Chardon *et al.* (2004) empirically suggested 5 times of the average EQF value per bin (5.35÷2× 5 = 13.38) as the threshold, which roughly corresponds to γ_10,0.05_ =13.32. Under γ_10,0.05_, there are 116 significant hotspots detected. The EQF threshold obtained by the *Q*-method is about 9.88 (corresponding to γ_177,0.05_, where 177 is the upper bound of *n*, *i.e., k* = 177), leading to the detection of 177 QTL hotspots, among which many of them are believed to be spurious since the EQF threshold obtained by the *Q*-method is too liberal (Neto *et al.* 2012; Yang *et al*. 2019). As compared to the result of Figure 4 in Yang *et al*. (2019) using trait grouping based on the general agronomic consideration (with 9 trait groups), the EQF thresholds obtained by the empirical trait grouping (with 71 trait groups) are higher, and the observed and expected hotspot numbers are closer to each other.

#### The yeast dataset

The yeast data consist of expression measurements on 5740 transcripts and 2956 genetic markers on 112 segregant strains (Brem and Kruglyak 2005). The expression measurements are further converted to normal quantiles for the hotspot analysis as described by Neto *et al.* (2012) and are available in the R package yeastqtl (https://github.com/byandell/qtlyeast). Among the 2956 markers, numerous markers are completely the same, and 1072 markers are different. The genome size is ~6345 cM. The average marker density is about one marker every ~5.92 cM. For comparison with the QTL hotspot detection analysis in Neto *et al*. (2012), the QTL mapping analysis was performed also using the regression interval mapping (Haley and Knott 1992) with the same bin size of 2 cM (Δ=2 cM) under a single-QTL model with the R/qtl software (Broman *et al.* 2003). The LOD scores at all bins for each trait were recorded to construct the LOD score matrix, and the 1.5-LOD 95% support interval (Neto *et al.* 2012) is used to construct the EQF matrix for the analyses that follow. The EQF-bin permutation with empirical trait grouping is performed to obtain the EQF thresholds.

For the same yeast data set, Neto *et al*. (2012) used a single-trait permutation LOD threshold of 3.45 corresponding to a 5% GWER to claim the QTL detection and construct the QTL matrix, and obtained 7.40 as the conservative LOD threshold (associated with hotspot size 1). Using the Gaussian process (Guo 2011), the relaxed LOD threshold for one trait ranges from ~3.05 to ~3.48, and the conservative LOD threshold for 5740 traits is ~6.17, respectively. We therefore use a sliding scale of empirical LOD thresholds of 3, 4, 5, 6, 7 and 8 for claiming QTL detection and constructing the six corresponding QTL matrices. Using the six LOD thresholds, among the 5740 molecular traits, there are 5586, 2797, 1740, 1232, 911 and 696 QTL detected for 3863, 2455, 1656, 1205, 895, and 688 traits, respectively, among which multiple QTL were detected for 1402, 325, 83, 27, 16, and 8 traits, respectively. As it should be, higher LOD thresholds will result in fewer detected QTL but with larger LOD scores. The QTL densities are less than one QTL per cM (~0.88, ~0.44, ~0.26, ~0.19, ~0.14 and ~0.11 under the six different LOD thresholds) for the six QTL matrices. The empirical trait grouping results in a total of 476, 562, 522, 451, 375 and 308 trait groups, respectively, under the six different LOD thresholds.

Figure 4, A-F, presents the 3- to 8-LOD EQF architectures of the yeast genome and the hotspots detected under different EQF thresholds (at 5% GWER). In Figure 4, A-F, the threshold values, γ_*n,α*_’s, for the test statistic qFreq(*n*)’s are coordinately represented by the left and right axes. For example, in the 3-LOD EQF architecture, the 1^st^ highest EQF peak is located in the bin [2,224] (223-225 cM of the 2^nd^ chromosome) with EQF value 168.68, which is not significant under the threshold γ_1,0.05_(3) = 169.68 for at least one spurious hotspot. Therefore, no hotspot is detected under γ_1,0.05_(3). The bin [2,224] is also the highest EQF peak with EQF values 119.47 and 71.52 in the 4- and 5-LOD EQF architectures, and is the 4^th^, 10^th^ and 14^th^ highest peak with EQF values 39.17, 17.74 and 8.46, respectively, in the 6-, 7- and 8-LOD EQF architectures. The γ_1,0.05_ thresholds in the 4- to 8-LOD EQF architectures are γ_1,0.05_(4) = 120.70, γ_1,0.05_(5) = 68.98, γ_1,0.05_(6) = 44.30, γ_1,0.05_(7) = 32.36 and γ_1,0.05_(8) = 25.72, respectively. Under these thresholds, the bin [2,224] is only significant in the 5-LOD EQF architectures. Using γ_1,0.05_, there is 0, 0, 1, 3, 2, 2 significant hotspots, respectively. The γ_2,0.05_ thresholds for detecting at least two hotspots are γ_2,0.05_(3) = 145.33, γ_2,0.05_(4) = 90.11, γ_2,0.05_(5) = 58.52, γ_2,0.05_(6) = 42.29, γ_2,0.05_(7) = 29.90 and γ_2,0.05_(8) = 24.35, respectively, in the six EQF architectures, and, under the six γ_2,0.05_ thresholds, there are 1, 2, 4, 3, 3 and 3 bins significant as hotspots (see more detailed chromosome-by-chromosome plots in Figure S3, 1-23) in practice. Similarly, the γ_5,0.05_ thresholds for detecting at least five hotspots are γ_5,0.05_(3) = 99.46, γ_5,0.05_(4) = 60.61, γ_5,0.05_(5) = 37.01, γ_5,0.05_(6) = 32.16, γ_5,0.05_(7) = 24.85, and γ_5,0.05_(8) = 19.47, respectively, and in practice there 4, 4, 8, 6, 5 and 5 bins significant as hotspots in the six EQF architecture. The EQF thresholds (*β*’s) obtained by the *Q*-method are 6.84, 5.15, 4.37, 3.77, 3.57 and 3.38 (corresponding to about γ_142,0.05_, γ_78,0.05_, γ_58,0.05_, γ_42,0.05_, γ_35,0.05_ and γ_28,0.05_), respectively, for the 3- to 8-LOD EQF architectures. Under the six *β*’s, there are a total of 137, 82, 48, 44, 32 and 27 significant hotspots, among which many of them are believed to be spurious since the *β*’s are too liberal due to ignoring the correlation structure among traits. In general, Figure 4, A-F, displays several obvious peaks in the six EQF architectures, indicating that there must exist several hotpots in the yeast genome.

Figure 5, A-F, presents the six EQF architectures of the bin [2,224] on the 2^nd^ chromosome. The bin [2,224] is the 1^st^ highest EQF bin with EQF values 168.68 (not significant γ_1,0.05_(3) = 169.68 but significant under γ_2,0.05_(3) = 145.33), 119.47 (not significant under γ_1,0.05_(4) = 120.70, but significant under γ_2,0.05_(4) = 90.11), 71.52 (significant under γ_1,0.05_(5) = 68.98) in the 3- to 5-LOD EQF architectures, respectively, and is the 4^th^, 10^th^ and 14^th^ highest peak with EQF values 39.17 (not significant under γ_2,0.05_(6) = 42.29, but significant under γ_3,0.05_(6) = 38.61), 17.74 (not significant under γ_7,0.05_(7) = 18.34, but significant under γ_8,0.05_(7) = 16.70) and 8.46 (not significant under γ_12,0.05_(8) = 8.75, but significant under γ_13,0.05_(8) = 8.32) in the 6- to 8-LOD EQF architectures. So, the top γ_*n,α*_ thresholds for the bin [2,224] are γ_2,0.05_(3), γ_2,0.05_(4), γ_1,0.05_(5), γ_3,0.05_(6), γ_8,0.05_(7) and γ_13,0.05_(8), with the values of *n* being 2, 2, 1, 3, 8 and 13, respectively, across the six EQF architectures, as outlined in Figure 5H, the top γ_*n,α*_ profile of the bin [2,224]. In Figure 5H, the values of *n* show an increasing pattern over the six LOD thresholds, showing that the bin [2, 224] is more significant as a hotspot by using the <6 LOD thresholds as compared to by using the ≥6 LOD thresholds. Therefore, the bin [2, 224] can be regarded as a major QTL hotspot containing relatively more QTL with <6 LOD scores and relatively less QTL with ≥ LOD scores (more QTL with moderate LOD scores and less QTL with large LOD scores. Hereinafter defining LOD scores <6 as moderate and LOD scores ≥6 as large LOD scores) as can be also observed from the distribution of LOD scores > 3 in Figure 5G.

**Figure 5.**
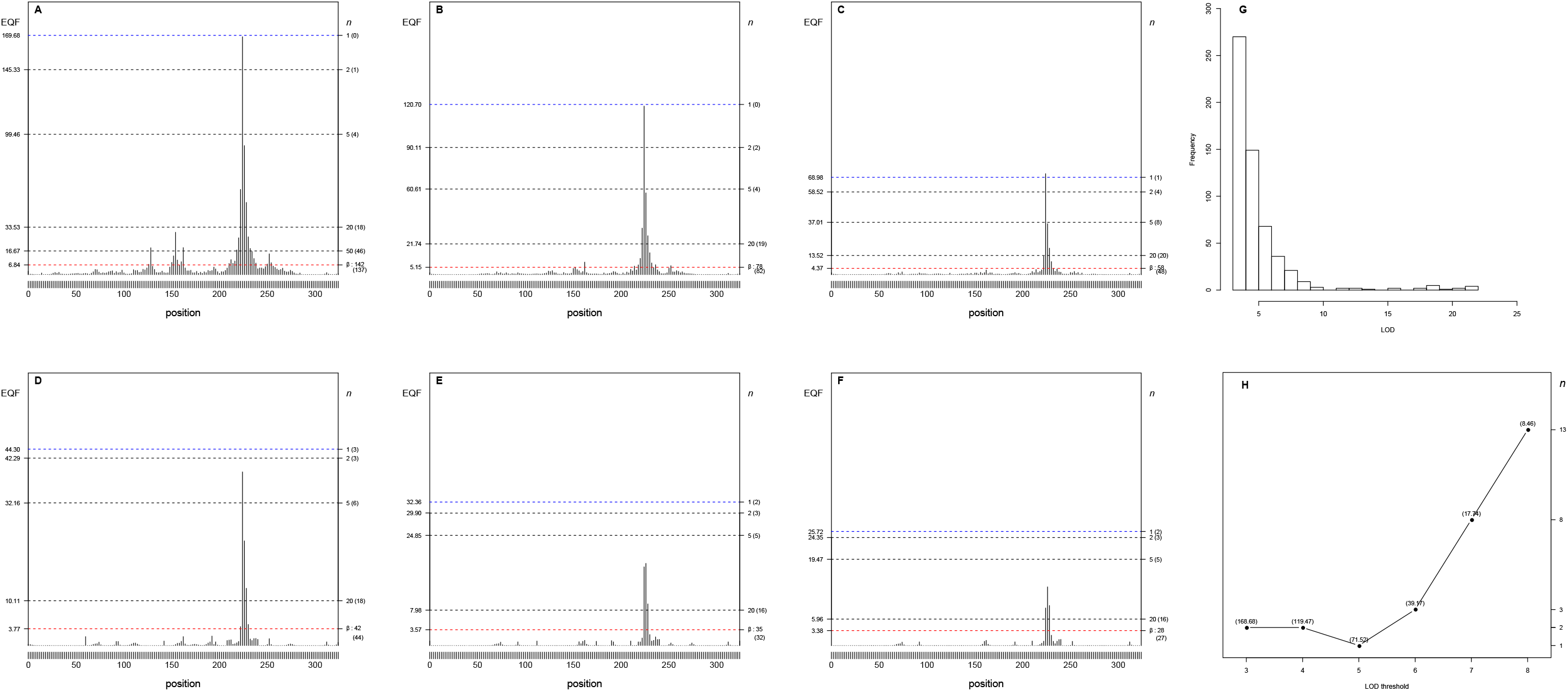
Panels (A-H) The 3- to 8-LOD EQF architectures of the 2^nd^ chromosome and the γ_*n*,0.05_ EQF thresholds at GWER of 5%. The left axis denotes the values of EQF, and the right axis denotes the values of *n*. (A-F) The peak at the bin [2,224] is significant as a hotspot under the γ_2,0.05_(3) = 145.33, γ_2,0.05_(4) = 90.11, γ_1,0.05_(5) = 68.98, γ_3,0.05_(6) = 38.61, γ_8,0.05_(7) = 16.70 and γ_13,0.05_(8) = 8.32 thresholds in the 3- to 8-LOD EQF architectures. The number in the bracket is the number of detected hotspots. (G) The distribution of LOD scores > 3 for the QTL at the bin [2,224]. (H) The top γ_*n*,0.05_ profile shows that the values of *n* are 2, 2, 1, 3, 8 and 13, showing an increasing pattern, across the 3- to 8-LOD EQF architectures. The number in the bracket is the EQF value of the bin.

The 2^nd^ highest peak in the 3-LOD EQF architecture is the bin [15,60]. The EQF values of the bin [15,60] are 144.27, 91.21, 62.39, 44.50, 29.94 and 18.18, respectively, in the 3- to 8-LOD EQF architectures (Figure 6A). The top γ_*n,α*_ thresholds are γ_3,0.05_(3) = 107.18, γ_2,0.05_(4) = 90.11, γ_2,0.05_(5) = 58.52, γ_1,0.05_(6) = 44.30, γ_2,0.05_(7) = 29.90 and γ_6,0.05_(8) = 17.85, with the values of *n* being 3, 2, 2, 1, 2 and 6, respectively, in the six EQF architectures (see Figure 6A bottom panel). Figure 6A depicts the top γ_*n,α*_ profile, showing that the values of *n* have an slightly increasing trend within the narrow range of 1 to 6 across the different EQF architectures. It tells that the bin [15,60] is a major hotspot containing QTL with balanced LOD-scores (moderate to large LOD scores) as can be also perceived in Figure 6A (upper panel). The 3^rd^ highest peak in the 3-LOD EQF architecture is the bin [14,242] (see Figure 4A). The EQF values of this bin are 106.16, 82.49, 64.66, 60.04, 43.55 and 32.20 in the 3-8 LOD EQF architectures (Figure 6B), and the top γ_*n,α*_ thresholds are γ_4,0.05_(3) = 101.26, γ_4,0.05_(4) = 81.14, γ_2,0.05_(5) = 58.52, γ_1,0.05_(6) = 44.30, γ_1,0.05_(7) = 32.36 and γ_1,0.05_(8) = 25.72, respectively (Figure 6B). Figure 6B (bottom panel) displays the top γ_*n,α*_ profile for the bin [14,242], showing that the values of *n* is flat within a narrow range from 1 to 4 over the EQF architectures. The flat pattern implies that the bin [14,242] is also a major hotspot containing the QTL with balanced LOD-scores. Figure 6B (upper panel) displays the distribution of LOD scores > 3 for the QTL in the bin [14,242]. The 24^th^ highest peak in the 3-LOD EQF architecture is the bin [3,89.5] (the first marker on chromosome 3 starting at the position of 9.5 cM). The EQF values of this bin are 29.58, 28.55, 24.72, 24.50, 25.35 and 22.54 in the 3- to 8-LOD EQF architectures (Figure 6C), the top γ_*n,α*_ thresholds are the top γ_*n,α*_ thresholds are γ_26,0.05_(3) = 29.46, γ_15,0.05_(4) = 28.30, γ_,0.05_(5) = 24.09, γ_8,0.05_(6) = 20.78, γ_5,0.05_(7) = 24.85 and γ_5,0.05_(8) = 19.47, respectively (see Figure 6C or Figure S3-4). Figure 6C (bottom panel) displays the top γ_*n,α*_ profile for the bin [3,89.5], showing a decreasing pattern with the value of *n* descending from 26 to 5 over the six thresholds. It means that the bin [3,89.5] is also a major hotspot containing relatively more QTL with ≥ 6 LOD scores. The distribution of LOD scores > 3 for the QTL at the bin [3,89.5] is presented in Figure 6C (upper panel).

**Figure 6.**
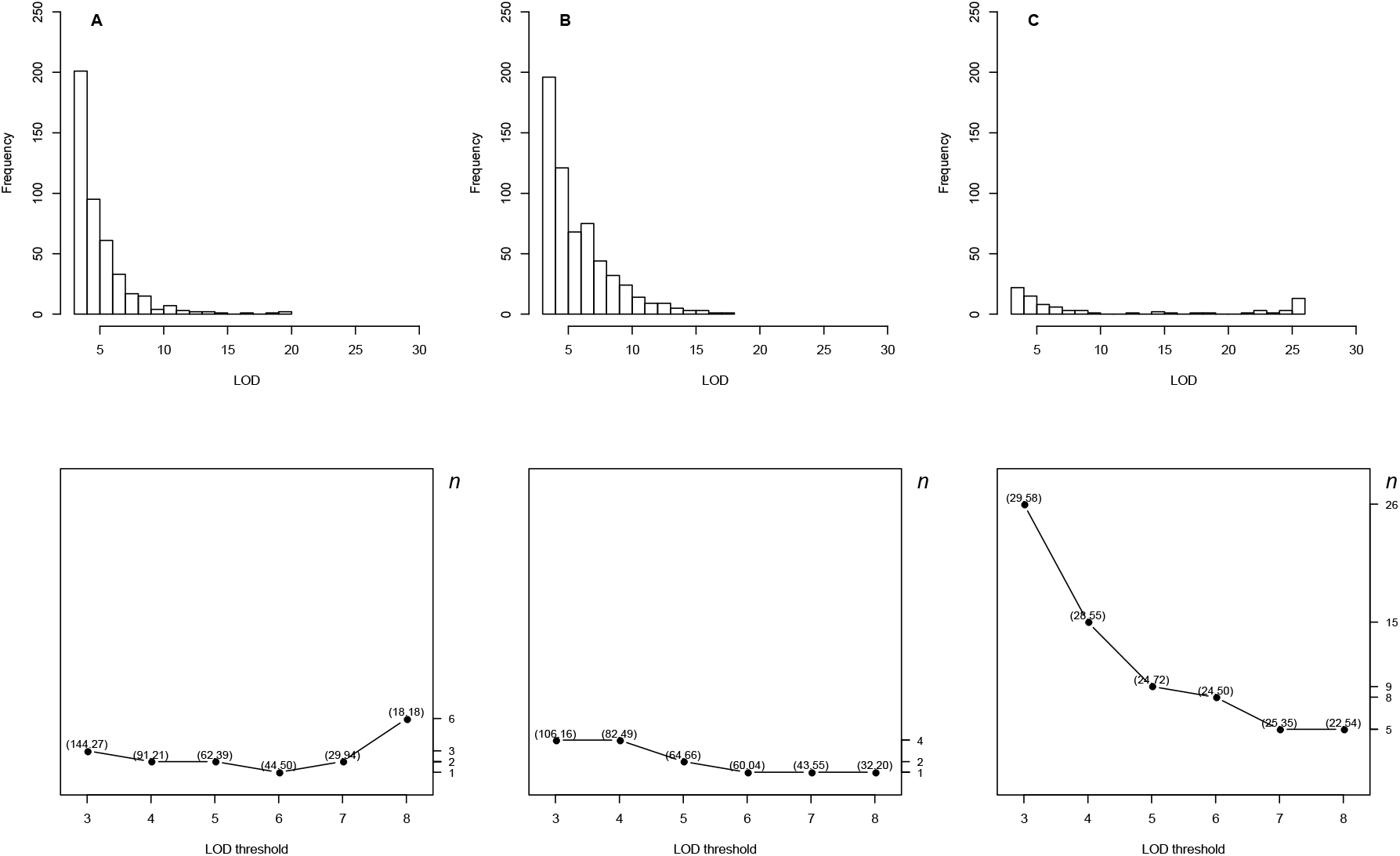
Panels (A-C)The distributions of QTL with LOD scores > 3 and the top γ_*n*,0.05_ profiles for the bins [15,60], [14,242] and [3,89.5] (the first marker on chromosome 3 starting at the position of 9.5 cM). The upper panels display the distributions of LOD scores, and the bottom panels show the top γ_*n*,0.05_ profiles. (A) The top γ_*n*,0.05_ profile displays that the *n* values have an increase trend within the narrow range from 1 to 6 over the 3- to 8-LOD thresholds, showing the bin [15,60] is a major hotspot containing QTL with balanced LOD-scores. (B) The top γ_*n*,0.05_ profile displays a flat pattern with the values of *n* varying within the range of from 1 to 4, showing the bin [14,242] is also a major hotspot containing the QTL with balanced LOD-scores. (C) The top γ_*n*,0.05_ profile has a decreasing trend over the 3- to 8-LOD thresholds, showing the bin [3,89.5] is a major hotspot is also a major hotspot containing relatively more QTL with ≥ 6 LOD scores. The number in the bracket is the EQF value of the bin.

In addition to the peak at [3,89.5], there is another peak at [3,57.5] on the 3^rd^ chromosome as shown in Figure 7, A-G. Figure 7H presents that the top γ_*n,α*_ profile of [3,57.5] has a flat pattern within the narrow range from 20 to 22 over the 3- to 8-LOD thresholds, showing that the bin [3,57.5] may be considered as a minor hotspot containing QTL with balanced LOD scores. It is also interesting to observe that the QTL at the bin [3,57.5] outnumber those at the bin [3,89.5] (see the panels for the LOD-score distributions of QTL in Figures 6C and 7G), but the EQF values are lower (except for the value in the 3-LOD architecture). It is because that the contributive QTL at the bin [3,57.5] have smaller LOD scores and tend to contribute less EQF values as compared to those at the bin [3,89.5]. The top γ_*n,α*_ profiles of the bins with local peaks and their histograms of LOD scores (> 3) in each of the 16 chromosomes in the 3- to 8-LOD EQF architectures are displayed in Figure S3, 1-23, (see supplementary material). In Figure S3-7, A-H, the EQF values of the bin [5,258] are 42.42, 12.15, 1.31, 0.08, 0.00164 and ~0, and the top γ_*n,α*_ thresholds are γ_17,0.05_(3) = 42.00, γ_31,0.05_(4) = 11.69, γ_230,0.05_(5) = 1.31, γ_1288,0.05_(6) = 0.08, γ_1483,0.05_(7) = 0.00157 and γ_1483,0.05_(8) ≈ 0, respectively, in the six EQF architectures. As the top γ_*n,α*_ profile displays an increasing trend from 17 to 1438 (Figure S3-7-H) implies that the bin [5,258] can be regarded as a secondary hotspot with very sparse QTL of strong LOD scores, which can be also observed in the histogram of LOD scores greater than 3 (Figure S3-7-G). The bin [5,258] will become less interesting with stricter LOD thresholds. In Figure S3-12, A-H, the EQF values of the bin [8,60] are 27.88, 23.81, 18.84, 15.61, 14.28 and 14.21, and the top γ_*n,α*_ thresholds are γ_28,0.05_(3) = 27.81, γ_18,0.05_(4) = 23.67, γ_14,0.05_(5) = 18.53, γ_11,0.05_(6) = 15.57, γ_10,0.05_(7) = 13.36and γ_8,0.05_(8) = 13.34, respectively (Figure S3-12-H). The top γ_*n,α*_ profile of bin [8,60] displays a decreasing trend in the values of *n* from 28 to 8 (Figure S3-12-G), showing that it consists of relatively more QTL with strong LOD scores and will be considered as a more significant hotspot using stricter LOD thresholds. In general, the significance status of a bin as a hotspot and the types of the hotspots can be well characterized by using the top γ_*n,α*_ profile.

**Figure 7.**
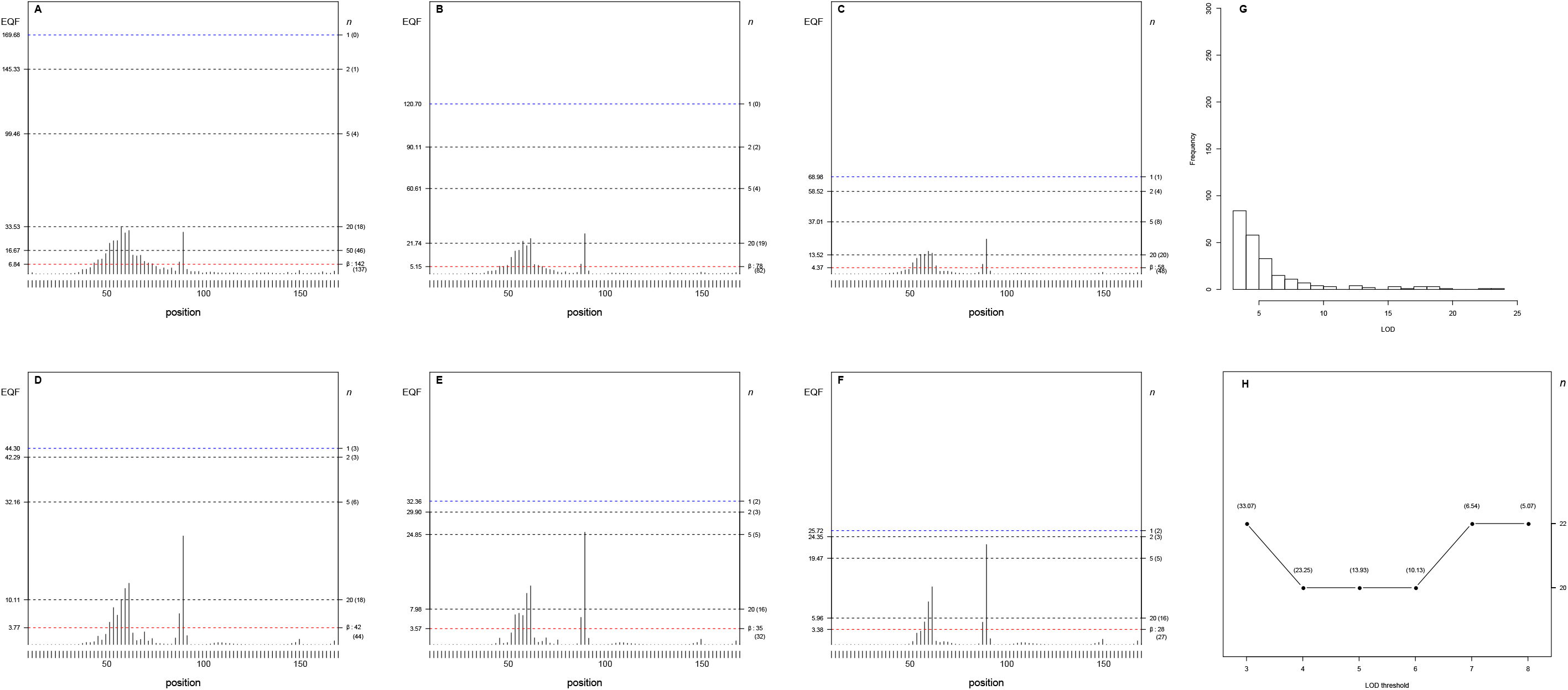
Panels (A-H) The 3− to 8-LOD EQF architectures of the 3^rd^ chromosome and the γ_*n*,0.05_ EQF thresholds at GWER of 5%. The left axis denotes the values of EQF, and the right axis denotes the values of *n*. (A-F) It shows the two peaks at the bins [3,57.5] and [3,89.5] are significant as the hotspots under the different γ_*n*,0.05_ EQF thresholds in the 3- to 8-LOD EQF architectures. The number in the bracket is the number of detected hotspots. (G) The distribution of LOD scores > 3 for the QTL at the bin [3,57.5]. (H) The top γ_*n*,0.05_ profile at the bin [3,57.5] shows that the values of *n* have a flat pattern varying from 20 to 22 across the 3- to 8-LOD EQF architectures. The number in the bracket is the EQF value of the bin. The LOD-score distribution of the QTL and the top γ_*n*,0.05_ profile at the bin [3,89.5] are shown in Figure 6C.

We also compared the above results obtained by the proposed statistical framework with those by the *Q*-, *N*-, and *NL*-methods presented in Neto *et al.* (2012). Using the EQF thresholds (*β*’s) of the *Q*-method (see Figure 4), there are 137, 82, 48, 44, 32 and 27 significant bins, among which most of them are believed to be spurious since the thresholds are known to be severely underestimated, in the six EQF architectures. The *N*-method detected five major hotspots on chromosomes 2, 3, 12, 14, and 15, and a suggestive hotspot (almost reaches significance) on chromosome 8, which are also detected by the *NL*-method (Neto *et al*. 2012) and by our statistical framework (correspond to the hotspots at the bins [2,224], [3,89.5], [8,60], [12,324], [14,242] and [15,60] in our analysis). Notably, our framework detects two hotspots at [3,61] and [3,89] on the 3^rd^ chromosome. Other small peaks on chromosomes 1, 4, 5, 7, 9, 13, and 16 considered by the *NL*-method are also detectable with our statistical framework (using less strict thresholds). Both the *NL*-method and our framework can assess the significance of hotspots with any type of LOD-score distribution. We follow Neto *et al*. to classify the hotspots into three types: (i) a hotspot composed of many QTL with moderate (<6) LOD, (ii) a hotspot consisting of a few QTL with strong (≧6) LOD scores, and (iii) a large hotspot containing QTL with a range of moderate to large LOD scores. In our statistical framework, the top γ_*n,α*_ profiles for the type i, ii and iii hotspots will respectively show increasing, decreasing and flat patterns. The *NL*-method summarized that the hotspots on chromosomes 2, 3, 12, 14, and 15 are of type iii, the hotspots on the 5^th^, 8^th^ and 13^th^ chromosomes is of type ii, and no hotspot is of type i. Using the top γ_*n,α*_ profiles, our framework concludes that, the hotspots at bin [2,224] and bin [5,258] on the 2^nd^ and 5^th^ chromosomes can be considered as being of type i (see Figure 5 and Figure S3-7), the hotspot at bin [3,89.5] on the 3^rd^ chromosome is of type ii (Figure 6C), not type iii by the *NL*-method, and the hotspots on the 12^th^, 14^th^ and 15^th^ chromosomes are of the same type (type iii) as the *NL*-method (see Figure 6 and Figure S3-16). Also, the hotspots at bin [8,60] on the 8^th^ chromosome and at bin [13, 516] on the 13^th^ chromosome are of the same type (type ii) as the *NL*-method (Figures S3-12 and S3-18). Neto *et al.* identified two significant peaks on each of the 2^nd^, 12^th^ and 15^th^ chromosomes; Nevertheless, we only observed a single significant peak on each of them, but detected two peaks (at bins [3,57.5] and [3,89.5]) on the 3^rd^ chromosome. The other small peaks at bins [4,464] and [7,30] on the 4^th^ and 7^th^ chromosomes are less interesting, but may be classified as type i according to their top γ_*n,α*_ profiles (see Figures S3-5 and S3-10). In general, the top γ_*n,α*_ profile can be used to characterize the three types of hotspots, and the results by the *NL-*method and our statistical framework are conformable in the detection and classification of QTL hotspots.

#### The pairwise phenotypic and genetic correlations in trait groups

Trait grouping intends to group the traits controlled by the same tightly linked or pleiotropic QTL together to cope with the correlation structure among the traits, avoiding the detection of spurious hotspots. We show by equations (2) and (3) and present in Table 1 that trait grouping based on the phenotypic or genetic correlations is not effective in combining the closely linked or pleiotropic traits. Also, the genetic correlations between monogenic pleiotropic traits are either −1 or +1, and the genetic correlations between multigenic pleiotropic traits can be arbitrary values between −1 and +1 (including zero), depending on the relative sizes and directions of effects and their linkage parameters. The above argument can be also justified by analyzing the pairwise phenotypic and genetic correlations among the traits (with closely linked or pleiotropic QTL) in the same trait group in the yeast data. Figure 8, A-F, displays the distributions for all pairwise phenotypic and genetic correlations among the traits in the first largest trait groups (the bin containing the most traits) in the 3- to 8-LOD EQF architectures. The largest trait groups contain 3123, 1411, 136, 78, 55 and 40 pleiotropic traits, among which there are 1297, 221, 5, 0, 0 and 0 traits are detected with multiple QTL, respectively, in the six EQF architectures. In the case of the 3-LOD EQF architecture (Figure 8A), as many (1297) pleiotropic traits are multigenic, their pairwise genetic correlations vary between +1 and −1, with a large proportion falling between −0.25 and 0.25. When the LOD thresholds become stricter, it will produce less detected QTL, less multigenic and more monogenic pleiotropic (or linked) traits (in proportion), causing that the pairwise genetic correlations of −1 or +1 gradually become the most common feature as shown in Figure 8, A-F (bottom panels). In Figure 8, D-F (bottom panels), the cases of the 6-LOD, 7-LOD and 8-LOD EQF architectures, all the 78, 55 and 40 traits are monogenic and hence the pairwise genetic correlations among them are either −1 or +1 (see subsection ***trait grouping***). The distributions for the pairwise phenotypic correlations among traits in the largest trait group are also presented in Figure 8, A-F (upper panels), showing the pleiotropic traits have a vaguer relationship with their phenotypic correlations. The distribution of pairwise phenotypic correlations shows a bell shape in the cases of 3- and 4-LOD EQF architectures (Figure 8, A-B upper panels), becomes a plateau type in the case of 4-LOD EQF architecture (Figure 8C upper panels), and has a bimodal pattern in the cases of 6-, 7- and 8-LOD EQF architectures (Figure 8, D-F upper panels), respectively. In general, the results in Figure 8 and the analytical derivations in equations (2) and (3) are compatible and confirm each other. The results also validate the use of the estimated QTL positions rather than the phenotypic or genetic correlations for trait grouping that intends to take into account the true linkages among QTL for avoiding the detection of spurious QTL hotspots in our statistical framework.

**Figure 8.**
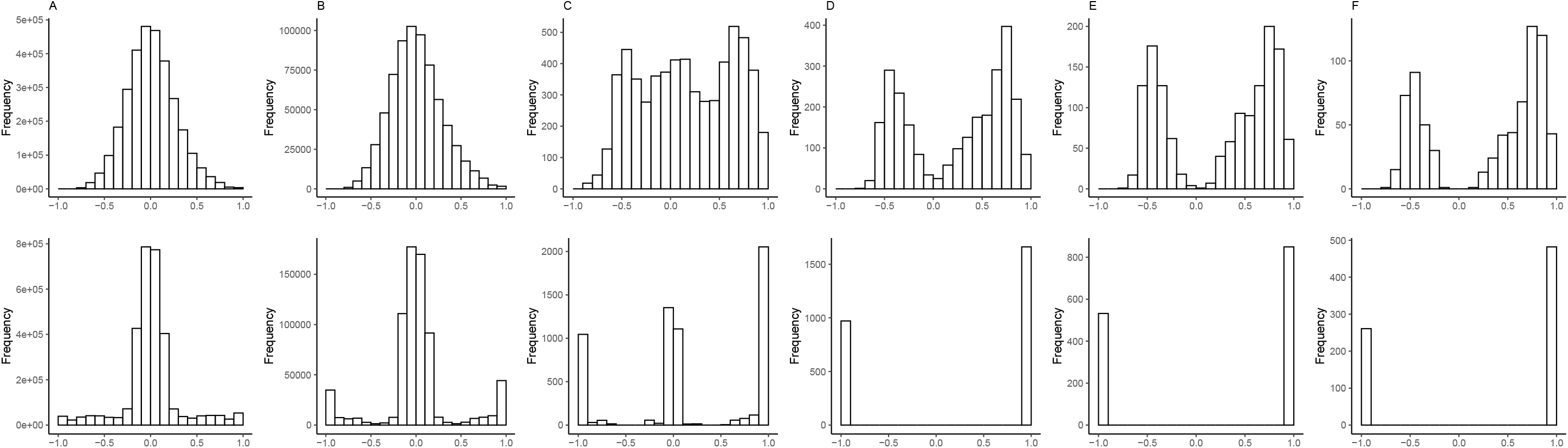
Panels (A-F) The distributions of pairwise phenotypic and genetic correlations in the first largest trait groups (containing most traits) of the 3- to 8-LOD EQF architectures. (A-F) The largest trait groups contain 3123, 1411, 136, 78, 55 and 40 pleiotropic traits (pleiotropic QTL), among which there are 1297, 221, 5, 0, 0 and 0 traits are detected with multiple QTL, respectively, in the six EQF architectures. The upper and bottom panels show the pairwise phenotypic and genetic correlations, respectively.

## DISCUSSION

Genome-wide detection of QTL hotspots requires first to collect the data with many QTL widespread in the genome, and then to construct the hotspot architectures and further to determine the thresholds for assessing the significance of QTL hotspots. The public databases and genetical genomics studies are two feasible ways to provide the data containing many QTL. The public databases curate abundant summarized QTL data for various traditional traits from numerous published studies (experiments), and the genetical genomics study can produce adequate individual-level data for many molecular traits in a single study for the detection of QTL hotspots. Using either type of data, statistical methods involved in determining the significance thresholds are of key importance and rely heavily on permutation analyses that perform on either the QTL matrix or individual-level data. The genetical genomics study allows to perform permutation analysis on both the QTL matrix and individual-level data, and the public database to date allows the analysis only on the QTL matrix to obtain the significance thresholds. Notably, permuting the QTL matrix has the outstanding advantage of very low computational cost, and permuting the individual-level data has the benefit of preserving the correlation structure among the traits but comes with very expensive computational cost (see **INTRODUCTION** section). Our statistical framework is deployed on the QTL matrices and hence can deal with both types of data with very low computational effort for QTL hotspot detection. Also, we introduce two special devices, trait grouping and top γ_*n,α*_ profile, into the framework, to address the concerns, including the correlation structure among traits and the magnitude of LOD scores within a hotspot, in the QTL hotspot detection. In trait grouping, by well using the QTL mapping results, the traits controlled by the same tightly linked and/or pleiotropic QTL are grouped and permuted together to obtain much stricter permutation thresholds. The trait grouping intends to directly take into account the true linkages among QTL, so as to cope with the correlation structure among the traits and dismiss spurious hotspots due to non-genetic correlations. The top γ_*n,α*_ profile is designed to outline the pattern of the top γ_*n,α*_ thresholds for a hotspot across the different EQF architectures constructed by using different LOD thresholds. If the top γ_*n,α*_ profile of a QTL hotspot shows an increasing (decreasing) pattern, it tells that the hotspot contains relatively fewer (more) QTL with strong LOD scores, as compare to other hotspots. A flat pattern of the top γ_*n,α*_ profile implies that the hotspot contains QTL with balanced LOD scores. Hence, the top γ_*n,α*_ profile can display the relative significance status of a hotspot at different EQF architectures, and can characterize and identify the types of QTL hotspots with different hotspot sizes (in terms of the EQF values) and LOD score distributions. In this way, our statistical framework can account for the correlation structure among the traits and identify the different types of hotspots with very low computational cost in the hotspot detection. Simulation study, numerical analysis and real examples are used to illustrate the proposed statistical framework, verify the related properties, and compare with the existing methods in the QTL hotspot detection.

It has been pointed out that the spurious hotspots may arise from the non-genetic correlations among traits or the use of liberal thresholds in the process of QTL hotspot detection (Darvasi 2003; Perez-Enciso 2004; Neto *et al.* 2012). The non-genetic correlations among traits are capable of inducing a spurious linkage, leading to excessive correlated traits being mapped to the similar regions and creating spurious QTL hotspots. The liberal thresholds arise out of ignoring the correlation structure among traits in the computation of the thresholds when assessing the significance of QTL hotspots (Breitling *et al.* 2008). Both imply the need to take the genetic correlations among traits as well as the true linkages between the underlying QTL into account for avoiding spurious hotspots in QTL hotspot detection. Linkages have been well known to be the main cause of genetic correlations among traits (Falconer and Mackey 1996). Such a fact is obvious for monogenic traits. However, for the multigenic traits, we show by equations (2) and (3) that they are not equivalent in the sense that strong linkages will not be necessary to create a significant genetic correlation between traits, with a value ranging from −1 to 1 including 0, simply because different linkage components may individually contribute positive or negative to the genetic correlation, but collectively combine to produce a low or no genetic correlation. This validates the approach to directly considering the true linkages instead of the genetic correlations in the analysis to dismiss spurious hotspots. The QTL mapping technique has proven to be powerful in estimating the QTL positions and related parameters to make inference about the true linkages (closely linked QTL or pleiotropic QTL) and to dissect the phenotypic correlation into genetic and non-genetic correlations for the traits (Jiang and Zeng 1995; Kao *et al.* 1999). By taking advantage of the QTL mapping results, our framework groups the traits with QTL being localized in the same bins (pleiotropic traits) together to take care of the true linkages, and permutes them together to compute much stricter hotspot size thresholds, so as to have the ability to control the genome-wide error rates and dismiss spurious hotspots due to non-genetic correlations and ignoring the correlation structure among traits. Instead of performing an infeasible multiple-trait joint analysis (Jiang and Zeng 1995) for testing the pleiotropy *vs.* close linkage among numerous QTL and traits, we directly treat the QTL being localized in the same bins as the tightly linked or pleiotropic QTL for trait grouping. If pleiotropic traits are partially grouped together in the computation, the thresholds are underestimated and the genome-wide error rates of falsely detecting a spurious hotspot will inflate, resulting in greater possibility of detecting spurious hotspots due using relaxed thresholds (Yang *et al.* 2019). Our statistical framework relied on the QTL detected by using appropriate QTL mapping methods. Here, the single-QTL regression interval mapping (Haley and Knott 1992) is adopted to estimate the QTL parameter. It would be also possible to extend our approach to using the EM interval mapping (Lander and Botstein 1989) and multiple-QTL interval mapping methods (Haley and Knott 1992; Kao *et al.* 1999; Sen and Churchill 2001). With multiple-QTL interval mapping, e.g., using the multiple-QTL mapping functions in QTL Cartographer or R/qtl (Basten *et al.* 1999 Broman *et al.* 2003), the LOD profile for each QTL adjusted for all other QTL can be obtained and used to construct the QTL matrices given some specified LOD thresholds. Once the QTL matrices are obtained, the subsequent steps are then straightforward to implement for hotspot detection.

The genetical genomics experiment is usually performed to produce transcript abundance of many genes (the transcription profile) at a time point or under a specific condition in the life stage of an organism. Then, by performing the QTL mapping analysis on the transcription profile followed by the analysis of QTL hotspot detection, we can obtain the EQF architecture (given a LOD threshold) to outline the QTL hotspot architecture along the genome for the organism. The QTL hotspot architecture actually summarizes the expressivity of genes at all the genomic positions and can be used to infer the networks among QTL hotspots, genes and traits at a time point for the organism (Yang et a. 2019). As the microarray technology for gene expressions becomes less expensive, it is possible to conduct the genetical genomics experiments at several time points during the life cycle of an organism to collect multiple transcription profiles for further investigating the behaviors of the QTL hotspots over the time course of the experiments. To take rice as an example, the genetical genomics experiments can be conducted at the vegetative, reproductive and ripening stages, or at multiple time points under the abiotic and biotic stresses (such as disease infection, pathogen attack, cold, drought, salt stresses) to obtain multiple transcription profiles. Then, we can perform the QTL mapping and hotspot detection analysis on the transcription profiles to obtain their respective QTL hotspot architectures at all the time points. Our statistical framework for QTL hotspot detection has a very low computational cost and hence is particular suitable for obtaining the QTL hotspot architectures for all the transcription profiles within a reasonable time frame as compared to the methods by permuting the individual-level data (without bothering the use of parallel computation on a cluster, see Neto *et al.* 2012). The collective QTL hotspot architectures can be used to discern the strengths (the pattern) of each specific QTL hotspot at different life stages or different time points after suffering the stresses. Through investigating the patterns of QTL hotspots across different time points, we can understand how the genes (genomic positions) express themselves differently at the different time points (stages) to outline their dynamic behaviors during the experiments. The study of together using the QTL public databases and the collective QTL hotspot architectures obtained from a series of genetic genomics experiments can help to explore the networks among the expressivity of genes, QTL hotspots and quantitative traits, as well as to provide deeper insight into the dynamic genomic activity for the organisms in broad areas of biological studies.

## Supplementary information

**Figure S1.**
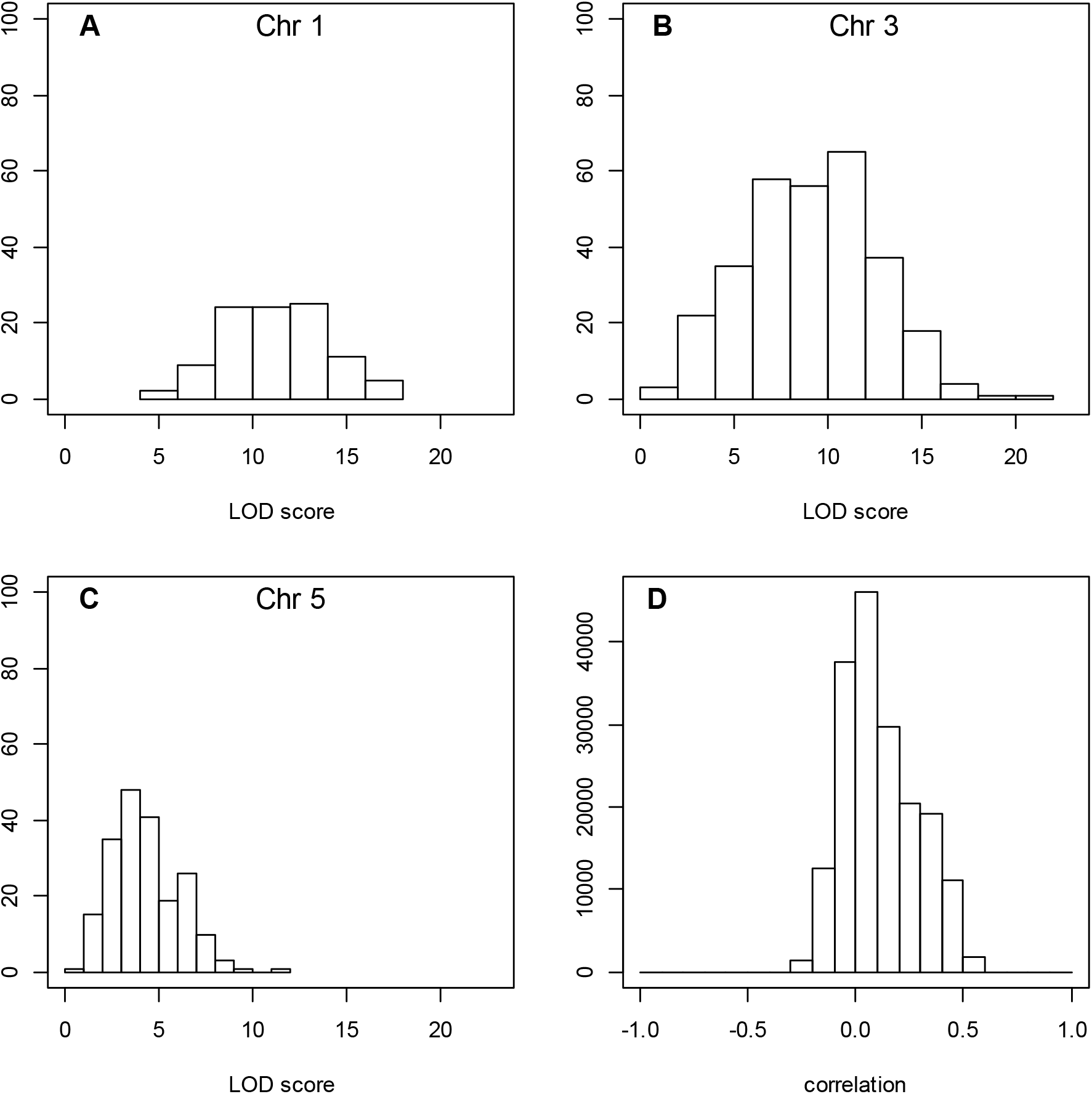
Panels (A-D) Distributions of hotspot LOD scores and pairwise correlations among traits for the simulated genetical genomics data. Panels (A), (B) and (C) show the LOD score distributions for the hotspot on chromosome 1, chromosome 3, and chromosome 5, respectively. The histograms show the distribution of the LOD scores of the traits composing the hotspot at the hotspot peak location. Panel (D) shows the distribution of the pairwise correlations among traits for the simulated data

**Figure S2.**
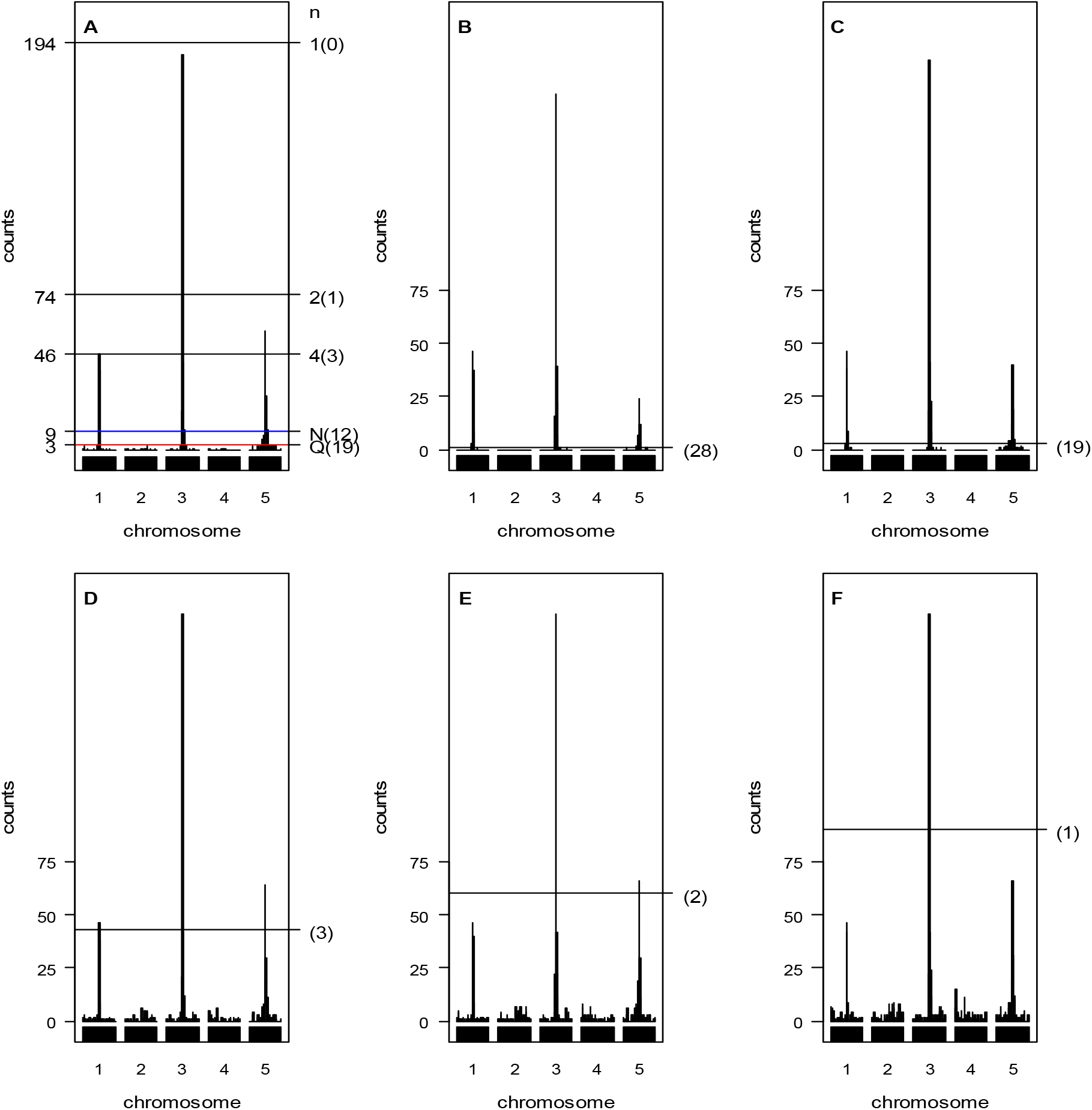
Panels (A-F) The proposed statistical procedure, *Q*-method, *N*-method and *NL*-method analyses for simulated example. Panels (A) Inferred hotspot architecture using a single-trait permutation LOD threshold of 2.47 corresponding to a GWER of 5% of falsely detecting at least one QTL somewhere in the genome. The hotspots on chromosomes 1, 3 and 5 have sizes 46, 188, and 57, respectively. The blue line at count 9 corresponds to the hotspot size threshold at a GWER of 5% according to the N- method. The red line at count 3 corresponds to the Q-method’s 5% significance threshold. The thresholds γ_1,0.05_, γ_2,0.05_, and γ_4,0.05_ obtained by the proposed procedure are 194, 74, and 46, respectively. Panels (B, C, D, E and F) Hotspot architecture inferred using different permutation thresholds by the NL-method; Hotspot architectures computed using QTL mapping LOD thresholds of 5.19 (B), 3.77 (C), 1.58 (D), 1.36 (E), and 1.13 (F) that aim to control GWER at a 5% error rate for spurious QTL hotspots of sizes 1, 3, 43, 60, and 90, respectively. The number in the bracket is the number of detected hotspots. Results are based on 1000 permutations. Q: The *Q-*method; N: The *N*-method.

**Figure S3-1.**
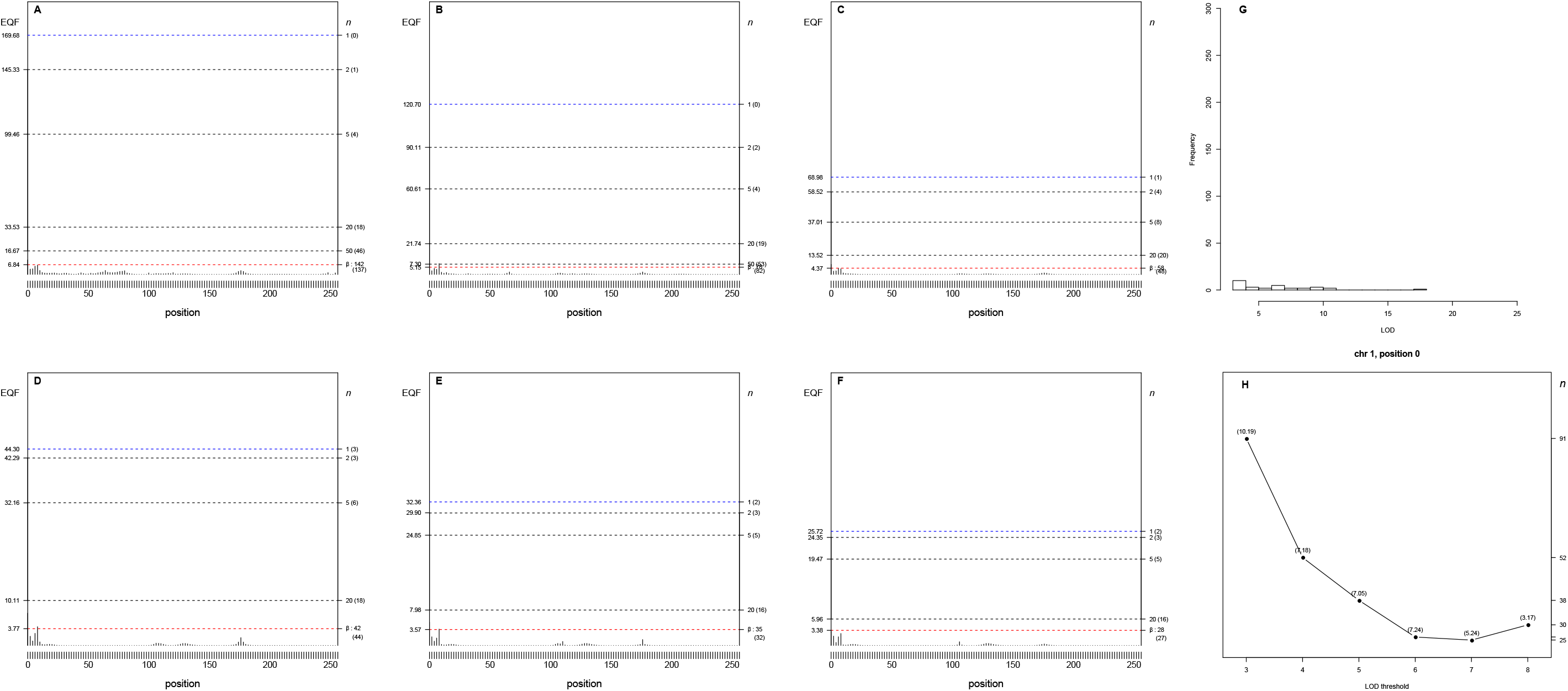
Panels (A-H) The 3- to 8-LOD EQF architectures of the 1^st^ chromosome and the γ_*n*,0.05_ EQF thresholds at GWER of 5%. The left axis denotes the values of EQF, and the right axis denotes the values of n. (A-F) It shows the one peak at the bin [1,0] is significant as the hotspots under the different γ_*n*,0.05_ EQF thresholds in the 3- to 8-LOD EQF architectures. The number in the bracket is the number of detected hotspots. (G) The distribution of LOD scores > 3 for the QTL at the bin [1,0]. (H) The top γ_*n*,0.05_ profile at the bin [1,0] shows that the values of n have a flat pattern varying from 91 to 25 across the 3- to 8-LOD EQF architectures. The number in the bracket is the EQF value of the bin.

**Figure S3-2.**
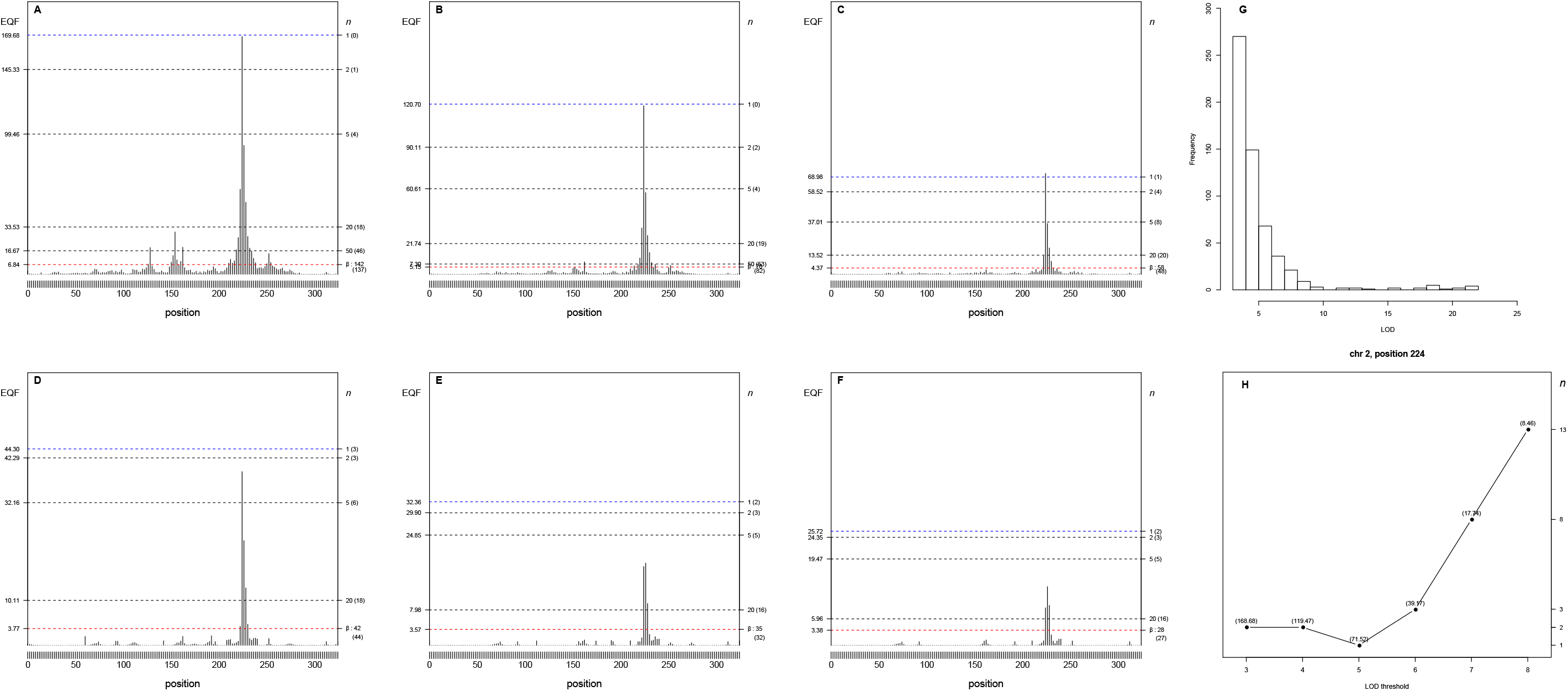
Panels (A-H) The 3− to 8-LOD EQF architectures of the 2^nd^ chromosome and the γ_*n*,0.05_ EQF thresholds at GWER of 5%. The left axis denotes the values of EQF, and the right axis denotes the values of n. (A-F) It shows the one peak at the bin [2,224] is significant as the hotspots under the different γ_*n*,0.05_ EQF thresholds in the 3- to 8-LOD EQF architectures. The number in the bracket is the number of detected hotspots. (G) The distribution of LOD scores > 3 for the QTL at the bin [2,224]. (H) The top γ_*n*,0.05_ profile at the bin [2,224] shows that the values of n have a flat pattern varying from 1 to 13 across the 3- to 8-LOD EQF architectures. The number in the bracket is the EQF value of the bin.

**Figure S3-3.**
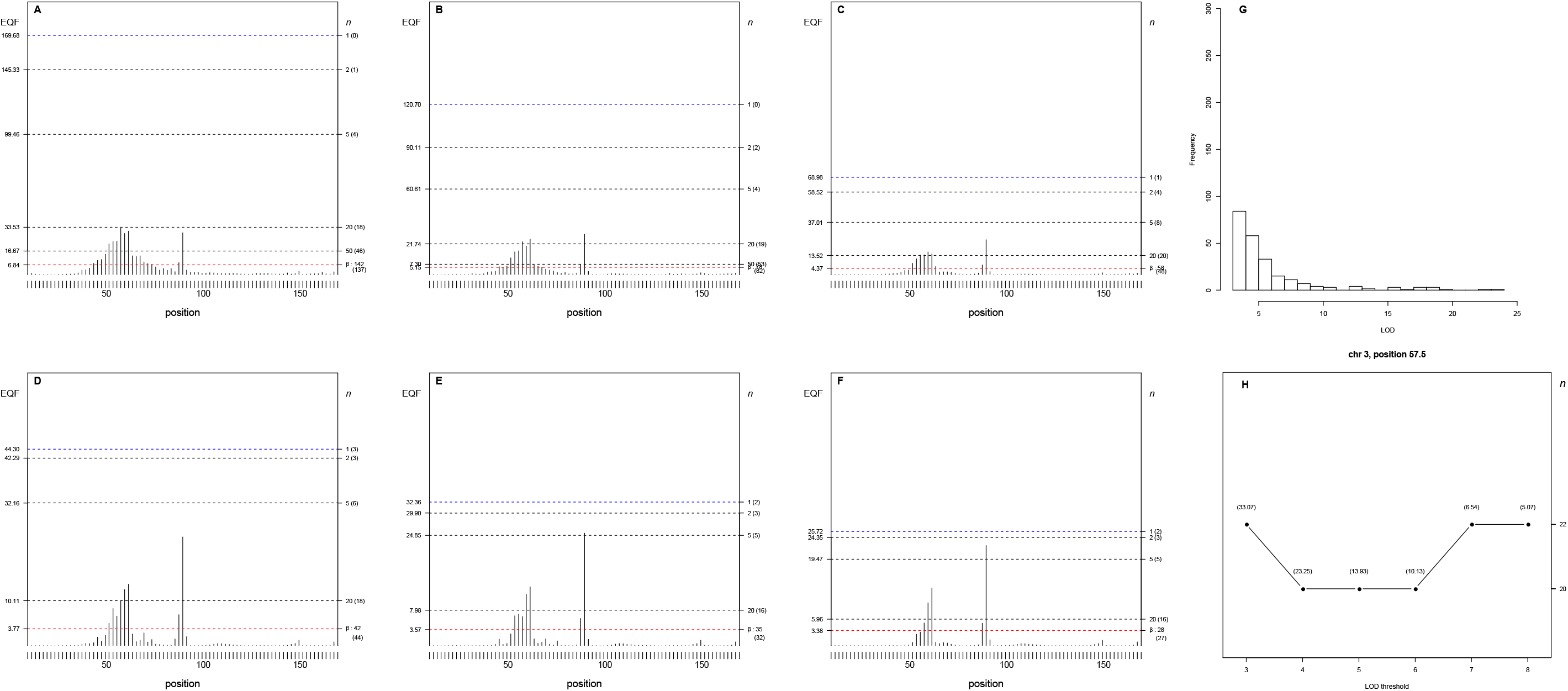
Panels (A-H) The 3- to 8-LOD EQF architectures of the 3^rd^ chromosome and the γ_*n*,0.05_ EQF thresholds at GWER of 5%. The left axis denotes the values of EQF, and the right axis denotes the values of n. (A-F) It shows the one peak at the bin [3,57.5] is significant as the hotspots under the different γ_*n*,0.05_ EQF thresholds in the 3- to 8-LOD EQF architectures. The number in the bracket is the number of detected hotspots. (G) The distribution of LOD scores > 3 for the QTL at the bin [3,57.5]. (H) The top γ_*n*,0.05_ profile at the bin [3,57.5] shows that the values of n have a flat pattern varying from 20 to 22 across the 3- to 8-LOD EQF architectures. The number in the bracket is the EQF value of the bin.

**Figure S3-4.**
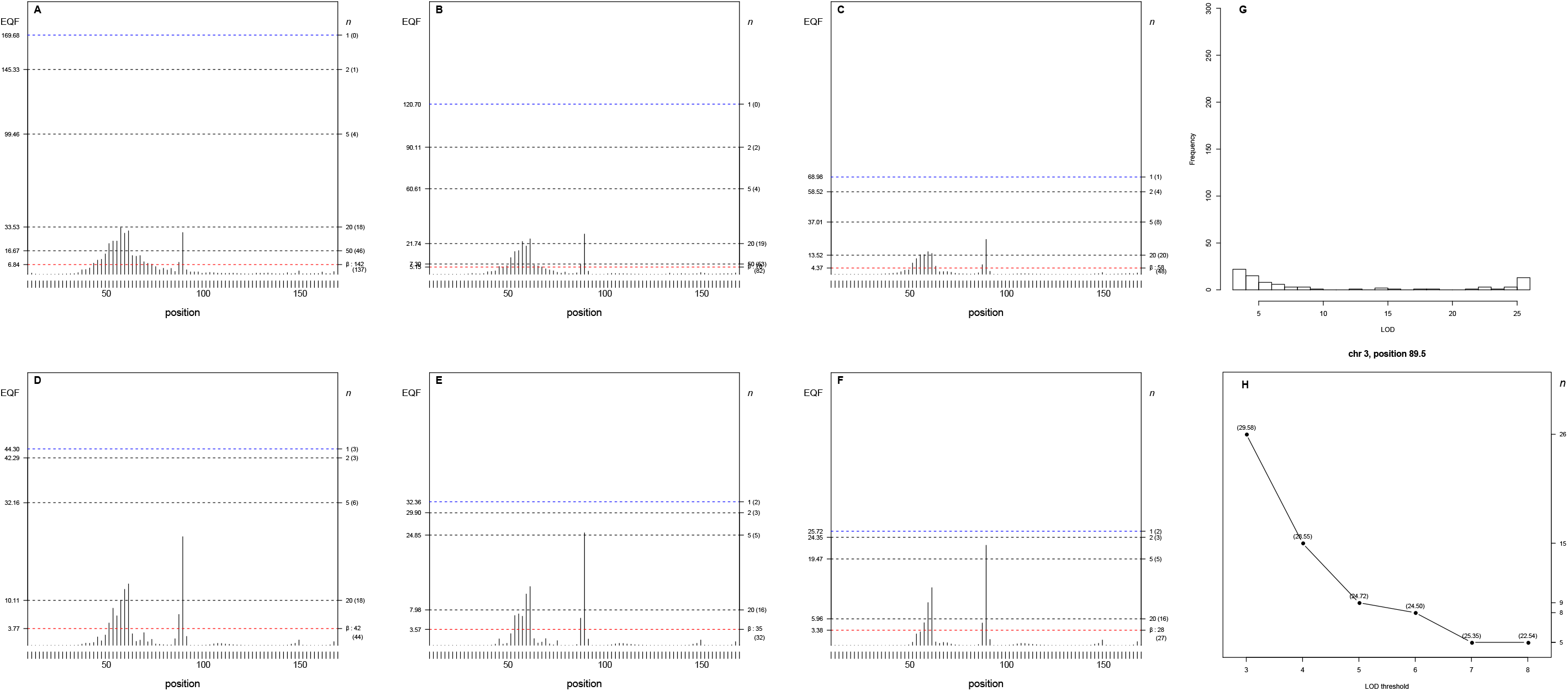
Panels (A-H) The 3- to 8-LOD EQF architectures of the 3^rd^ chromosome and the γ_*n*,0.05_ EQF thresholds at GWER of 5%. The left axis denotes the values of EQF, and the right axis denotes the values of n. (A-F) It shows the one peak at the bin [3,89.5] is significant as the hotspots under the different γ_*n*,0.05_ EQF thresholds in the 3- to 8-LOD EQF architectures. The number in the bracket is the number of detected hotspots. (G) The distribution of LOD scores > 3 for the QTL at the bin [3,89.5]. (H) The top γ_*n*,0.05_ profile at the bin [3,89.5] shows that the values of n have a flat pattern varying from 26 to 5 across the 3- to 8-LOD EQF architectures. The number in the bracket is the EQF value of the bin.

**Figure S3-5.**
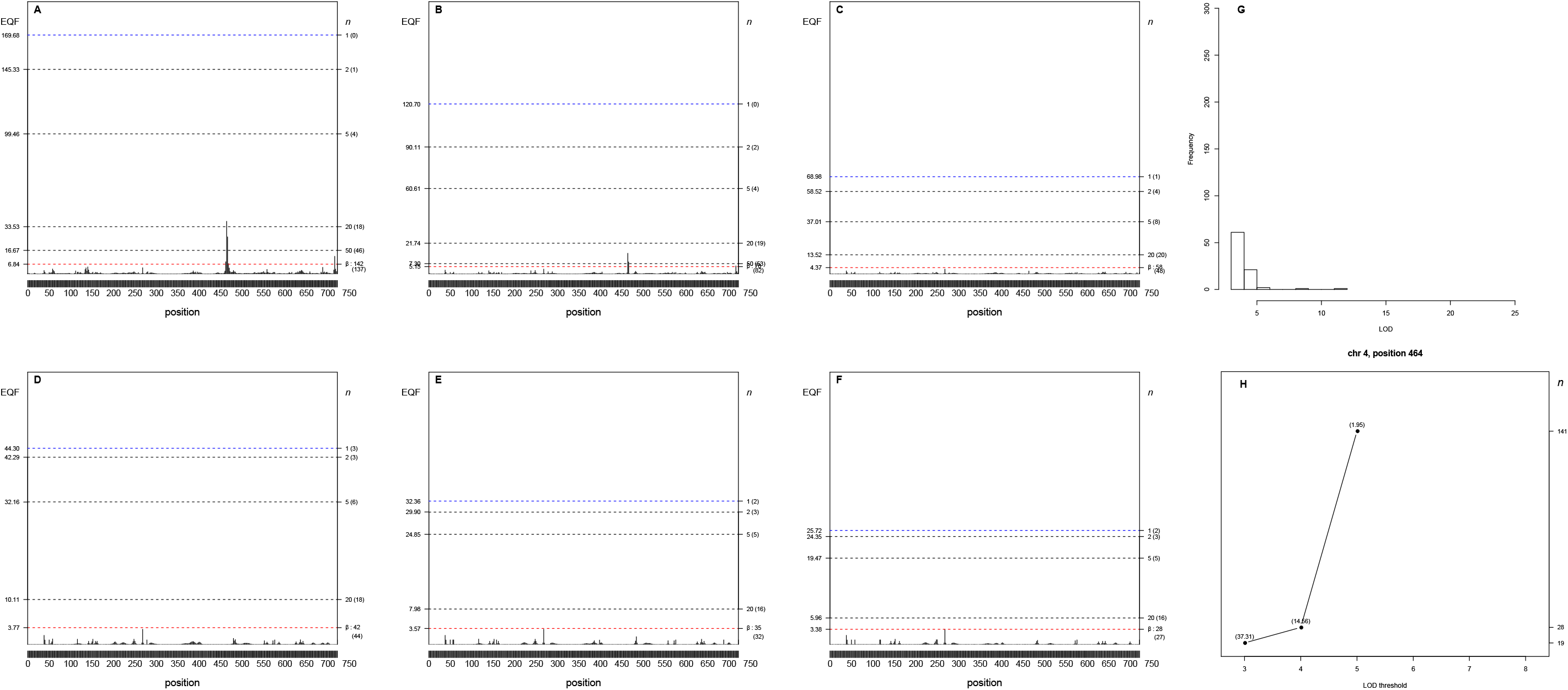
Panels (A-H) The 3- to 8-LOD EQF architectures of the 4^th^ chromosome and the γ_*n*,0.05_ EQF thresholds at GWER of 5%. The left axis denotes the values of EQF, and the right axis denotes the values of n. (A-F) It shows the one peak at the bin [4,464] is significant as the hotspots under the different γ_*n*,0.05_ EQF thresholds in the 3- to 8-LOD EQF architectures. The number in the bracket is the number of detected hotspots. (G) The distribution of LOD scores > 3 for the QTL at the bin [4,464]. (H) The top γ_*n*,0.05_ profile at the bin [4,464] shows that the values of n have a flat pattern varying from 19 to 141 across the 3- to 8-LOD EQF architectures. The number in the bracket is the EQF value of the bin.

**Figure S3-6.**
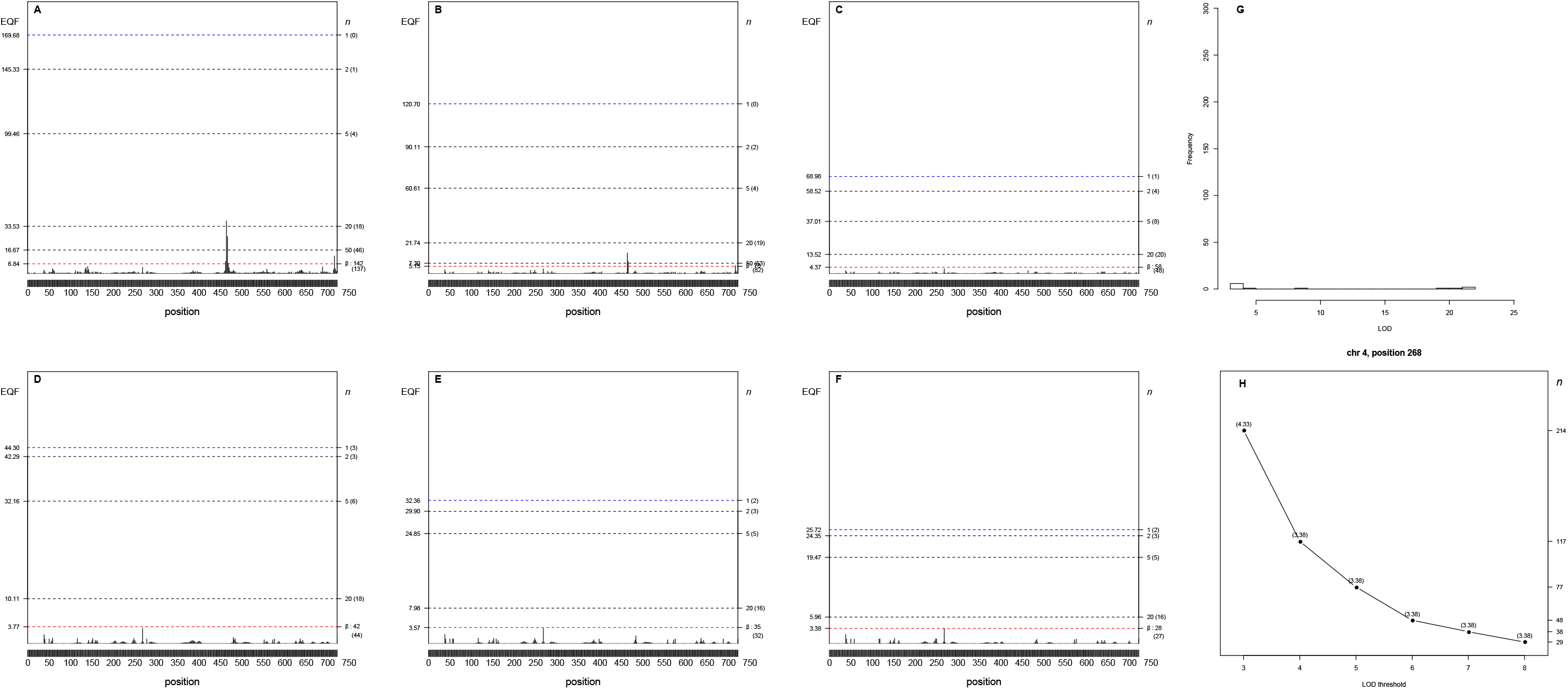
Panels (A-H) The 3- to 8-LOD EQF architectures of the 4^th^ chromosome and the γ_*n*,0.05_ EQF thresholds at GWER of 5%. The left axis denotes the values of EQF, and the right axis denotes the values of n. (A-F) It shows the one peak at the bin [4, 268] is significant as the hotspots under the different γ_*n*,0.05_ EQF thresholds in the 3- to 8-LOD EQF architectures. The number in the bracket is the number of detected hotspots. (G) The distribution of LOD scores > 3 for the QTL at the bin [4,268]. (H) The top γ_*n*,0.05_ profile at the bin [4, 268] shows that the values of n have a flat pattern varying from 214 to 29 across the 3- to 8-LOD EQF architectures. The number in the bracket is the EQF value of the bin.

**Figure S3-7.**
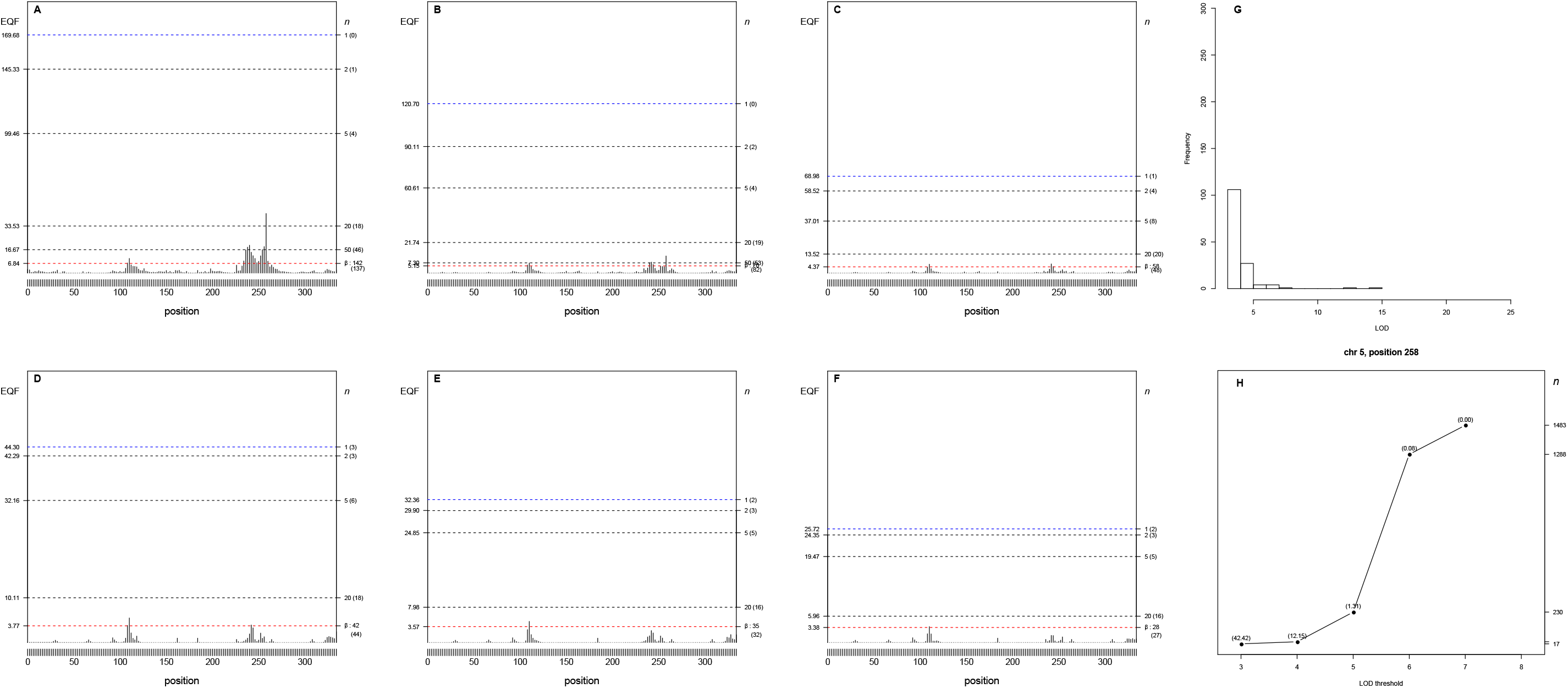
Panels (A-H) The 3- to 8-LOD EQF architectures of the 5^th^ chromosome and the γ_*n*,0.05_ EQF thresholds at GWER of 5%. The left axis denotes the values of EQF, and the right axis denotes the values of n. (A-F) It shows the one peak at the bin [5, 258] is significant as the hotspots under the different γ_*n*,0.05_ EQF thresholds in the 3- to 8-LOD EQF architectures. The number in the bracket is the number of detected hotspots. (G) The distribution of LOD scores > 3 for the QTL at the bin [5,258]. (H) The top γ_*n*,0.05_ profile at the bin [5, 258] shows that the values of n have a flat pattern varying from 17 to 1483 across the 3- to 8-LOD EQF architectures. The number in the bracket is the EQF value of the bin.

**Figure S3-8.**
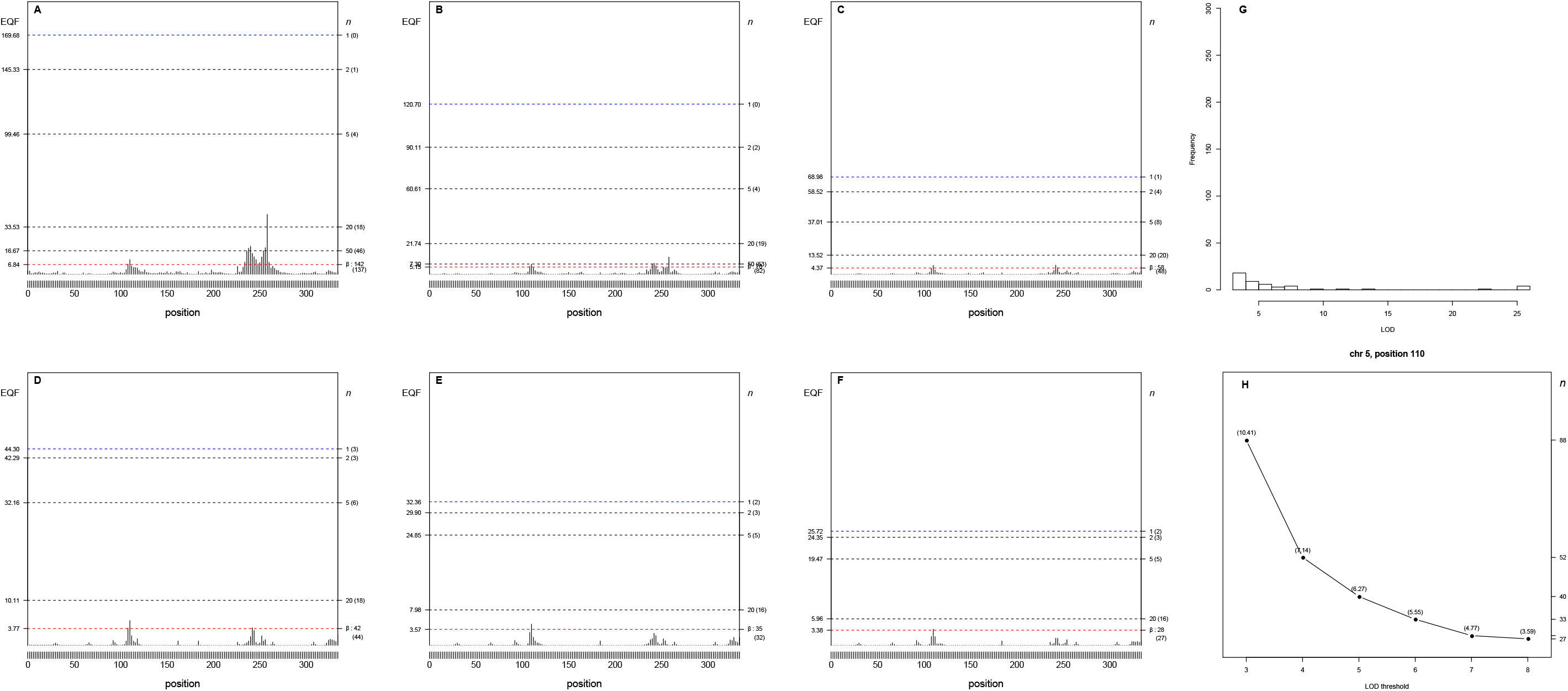
Panels (A-H) The 3- to 8-LOD EQF architectures of the 5^th^ chromosome and the γ_*n*,0.05_ EQF thresholds at GWER of 5%. The left axis denotes the values of EQF, and the right axis denotes the values of n. (A-F) It shows the one peak at the bin [5, 110] is significant as the hotspots under the different γ_*n*,0.05_ EQF thresholds in the 3- to 8-LOD EQF architectures. The number in the bracket is the number of detected hotspots. (G) The distribution of LOD scores > 3 for the QTL at the bin [5,110]. (H) The top γ_*n*,0.05_ profile at the bin [5, 110] shows that the values of n have a flat pattern varying from 88 to 27 across the 3- to 8-LOD EQF architectures. The number in the bracket is the EQF value of the bin.

**Figure S3-9.**
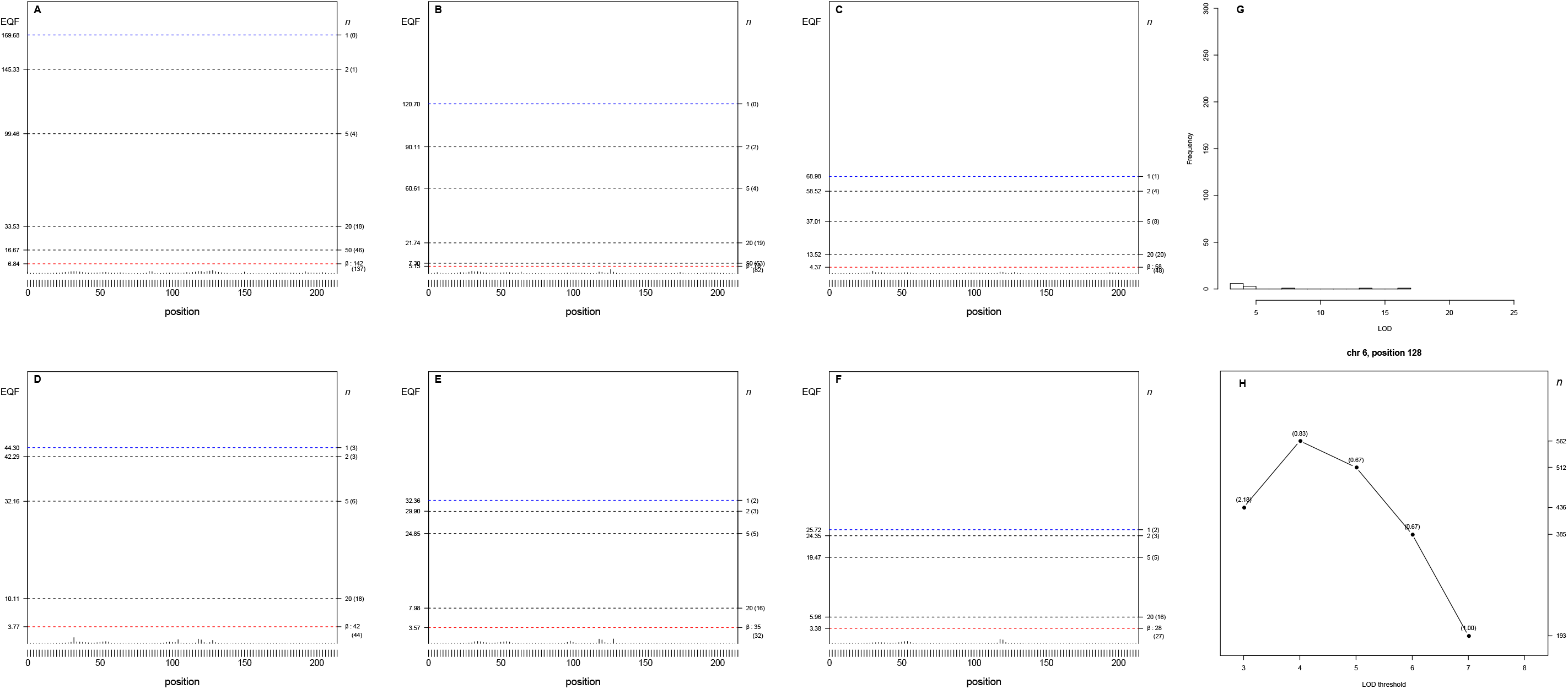
Panels (A-H) The 3- to 8-LOD EQF architectures of the 6^th^ chromosome and the γ_*n*,0.05_ EQF thresholds at GWER of 5%. The left axis denotes the values of EQF, and the right axis denotes the values of n. (A-F) It shows the one peak at the bin [6,128] is significant as the hotspots under the different γ_*n*,0.05_ EQF thresholds in the 3- to 8-LOD EQF architectures. The number in the bracket is the number of detected hotspots. (G) The distribution of LOD scores > 3 for the QTL at the bin [6,128]. (H) The top γ_*n*,0.05_ profile at the bin [6,128] shows that the values of n have a flat pattern varying from 562 to 193 across the 3- to 8-LOD EQF architectures. The number in the bracket is the EQF value of the bin.

**Figure S3-10.**
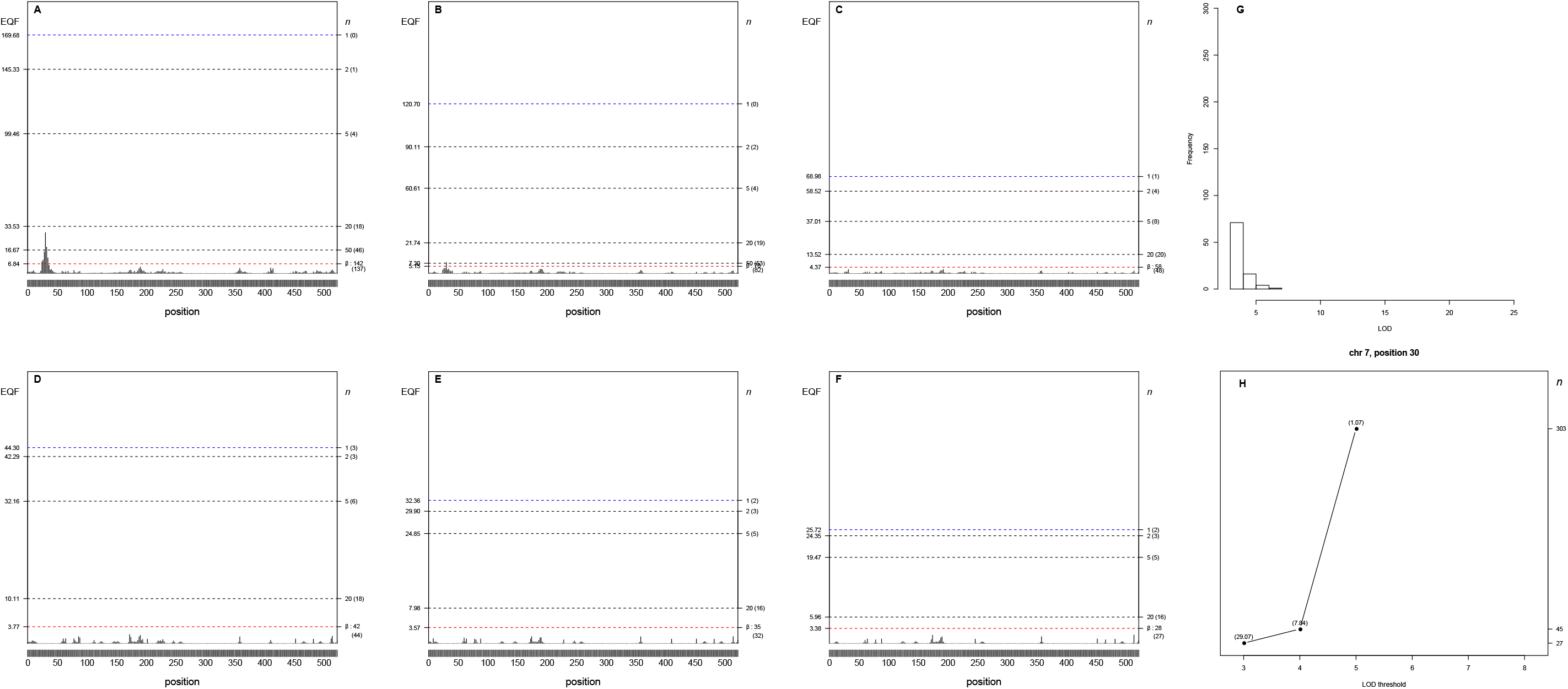
Panels (A-H) The 3- to 8-LOD EQF architectures of the 7^th^ chromosome and the γ_*n*,0.05_ EQF thresholds at GWER of 5%. The left axis denotes the values of EQF, and the right axis denotes the values of n. (A-F) It shows the one peak at the bin [7,30] is significant as the hotspots under the different γ_*n*,0.05_ EQF thresholds in the 3- to 8-LOD EQF architectures. The number in the bracket is the number of detected hotspots. (G) The distribution of LOD scores > 3 for the QTL at the bin [7,30]. (H) The top γ_*n*,0.05_ profile at the bin [7,30] shows that the values of n have a flat pattern varying from 27 to 303 across the 3- to 8-LOD EQF architectures. The number in the bracket is the EQF value of the bin.

**Figure S3-11.**
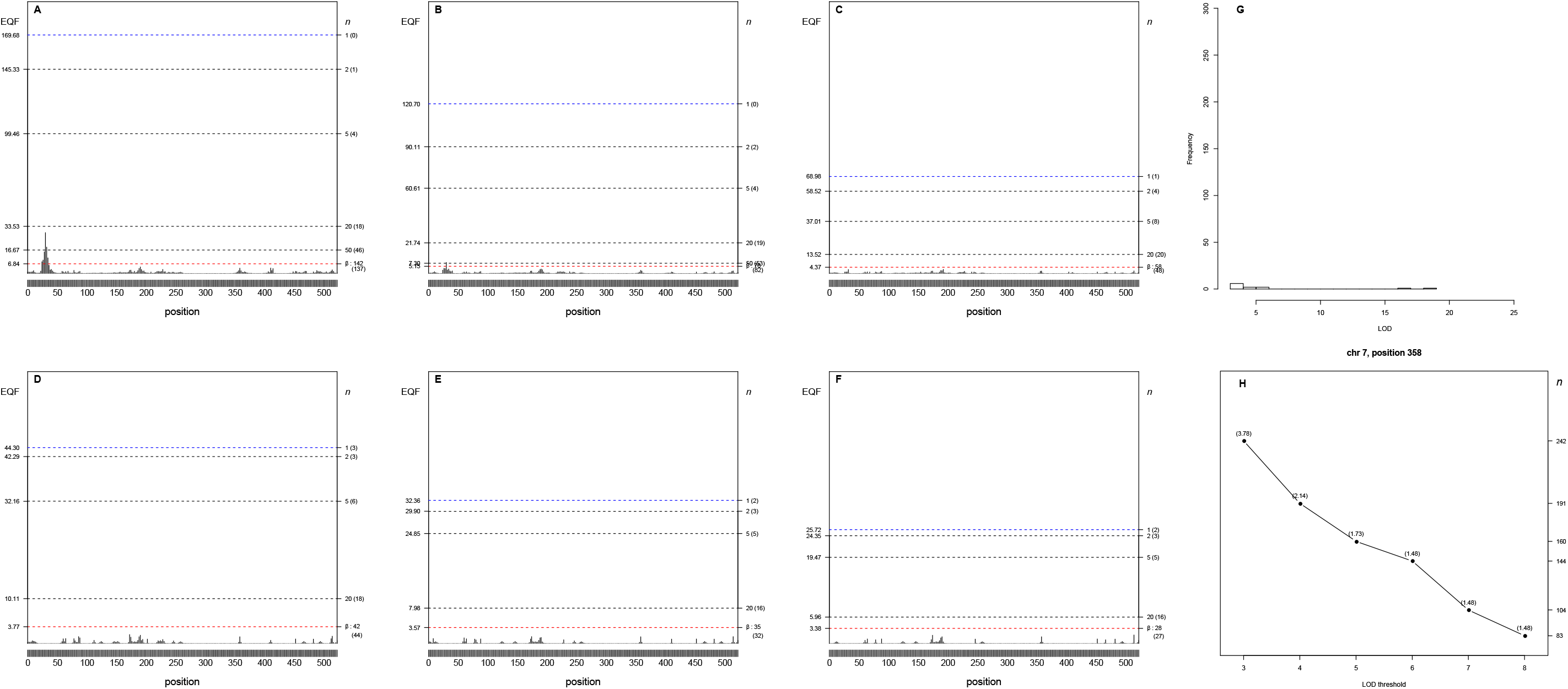
Panels (A-H) The 3- to 8-LOD EQF architectures of the 7^th^ chromosome and the γ_*n*,0.05_ EQF thresholds at GWER of 5%. The left axis denotes the values of EQF, and the right axis denotes the values of n. (A-F) It shows the one peak at the bin [7,358] is significant as the hotspots under the different γ_*n*,0.05_ EQF thresholds in the 3- to 8-LOD EQF architectures. The number in the bracket is the number of detected hotspots. (G) The distribution of LOD scores > 3 for the QTL at the bin [7,358]. (H) The top γ_*n*,0.05_ profile at the bin [7, 358] shows that the values of n have a flat pattern varying from 242 to 83 across the 3- to 8-LOD EQF architectures. The number in the bracket is the EQF value of the bin.

**Figure S3-12.**
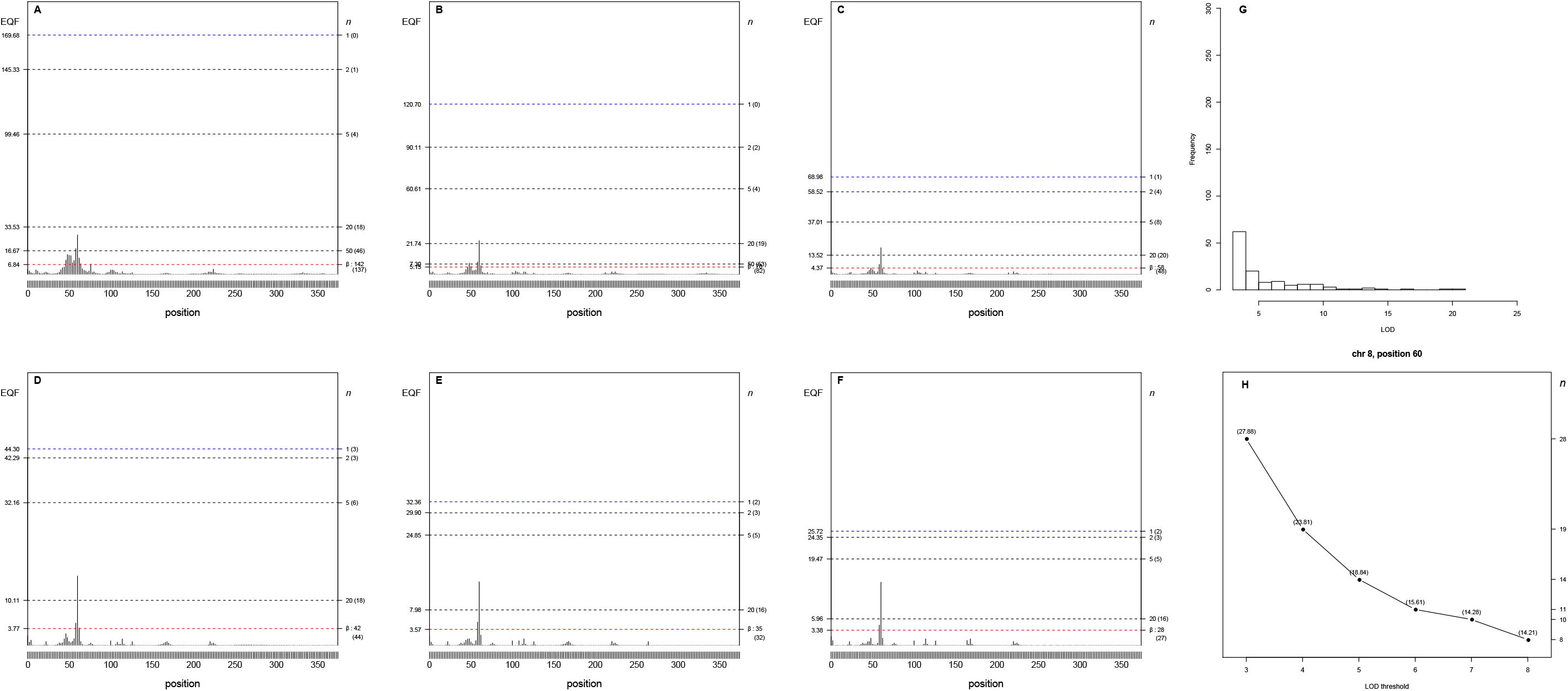
Panels (A-H) The 3- to 8-LOD EQF architectures of the 8^th^ chromosome and the γ_*n*,0.05_ EQF thresholds at GWER of 5%. The left axis denotes the values of EQF, and the right axis denotes the values of n. (A-F) It shows the one peak at the bin [8,60] is significant as the hotspots under the different γ_*n*,0.05_ EQF thresholds in the 3- to 8-LOD EQF architectures. The number in the bracket is the number of detected hotspots. (G) The distribution of LOD scores > 3 for the QTL at the bin [8,60]. (H) The top γ_*n*,0.05_ profile at the bin [8,60] shows that the values of n have a flat pattern varying from 28 to 8 across the 3- to 8-LOD EQF architectures. The number in the bracket is the EQF value of the bin.

**Figure S3-13.**
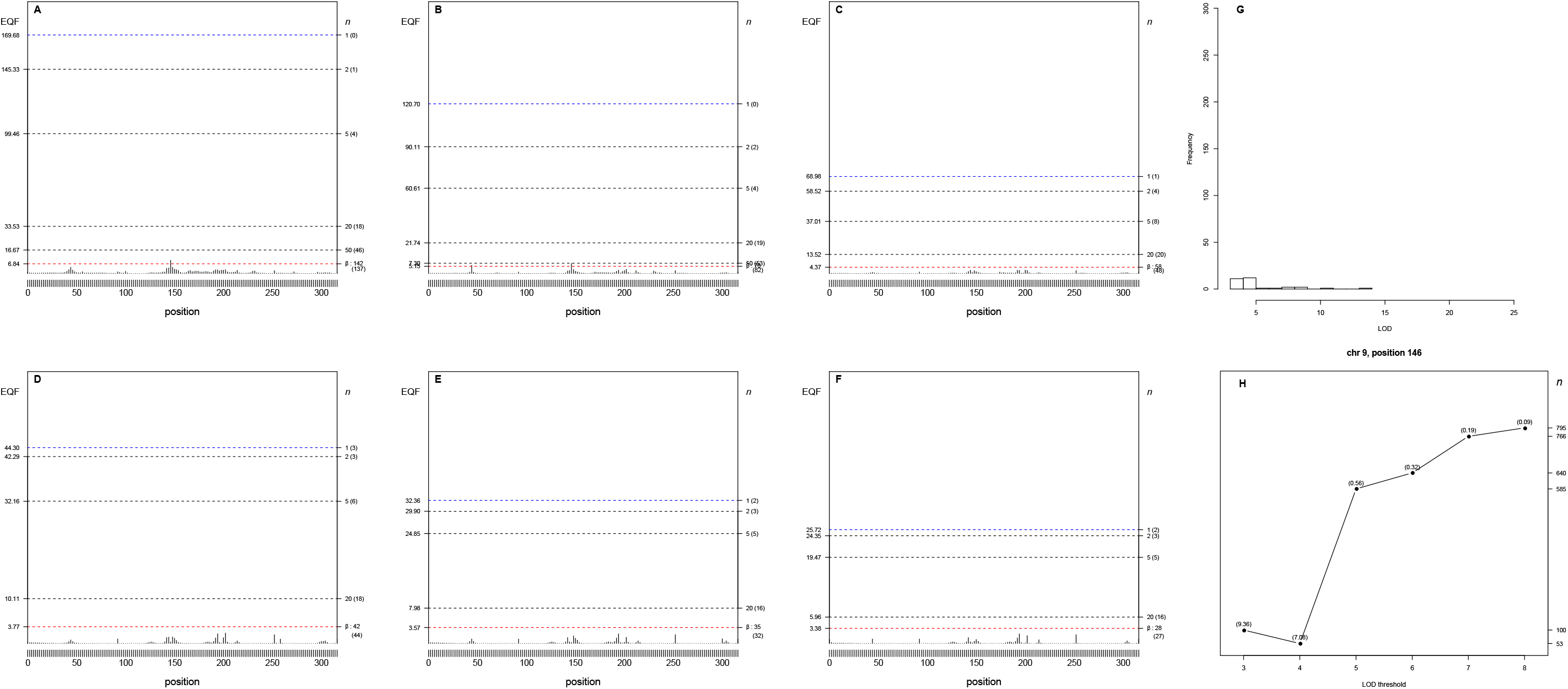
Panels (A-H) The 3- to 8-LOD EQF architectures of the 9^th^ chromosome and the γ_*n*,0.05_ EQF thresholds at GWER of 5%. The left axis denotes the values of EQF, and the right axis denotes the values of n. (A-F) It shows the one peak at the bin [9,146] is significant as the hotspots under the different γ_*n*,0.05_ EQF thresholds in the 3- to 8-LOD EQF architectures. The number in the bracket is the number of detected hotspots. (G) The distribution of LOD scores > 3 for the QTL at the bin [9,146]. (H) The top γ_*n*,0.05_ profile at the bin [9,146] shows that the values of n have a flat pattern varying from 53 to 975 across the 3- to 8-LOD EQF architectures. The number in the bracket is the EQF value of the bin.

**Figure S3-14.**
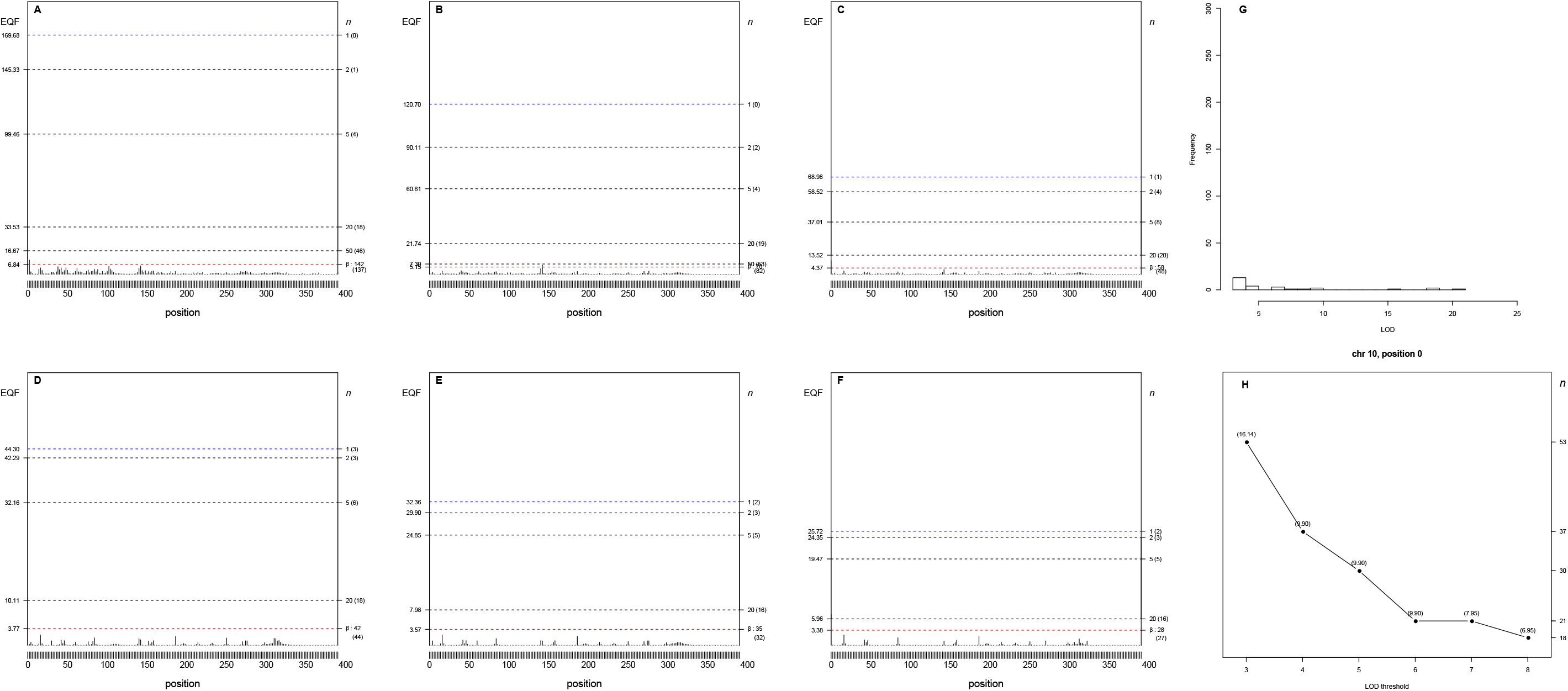
Panels (A-H) The 3- to 8-LOD EQF architectures of the 10^th^ chromosome and the γ_*n*,0.05_ EQF thresholds at GWER of 5%. The left axis denotes the values of EQF, and the right axis denotes the values of n. (A-F) It shows the one peak at the bin [10,0] is significant as the hotspots under the different γ_*n*,0.05_ EQF thresholds in the 3- to 8-LOD EQF architectures. The number in the bracket is the number of detected hotspots. (G) The distribution of LOD scores > 3 for the QTL at the bin [10,0]. (H) The top γ_*n*,0.05_ profile at the bin [10,0] shows that the values of n have a flat pattern varying from 53 to 18 across the 3- to 8-LOD EQF architectures. The number in the bracket is the EQF value of the bin.

**Figure S3-15.**
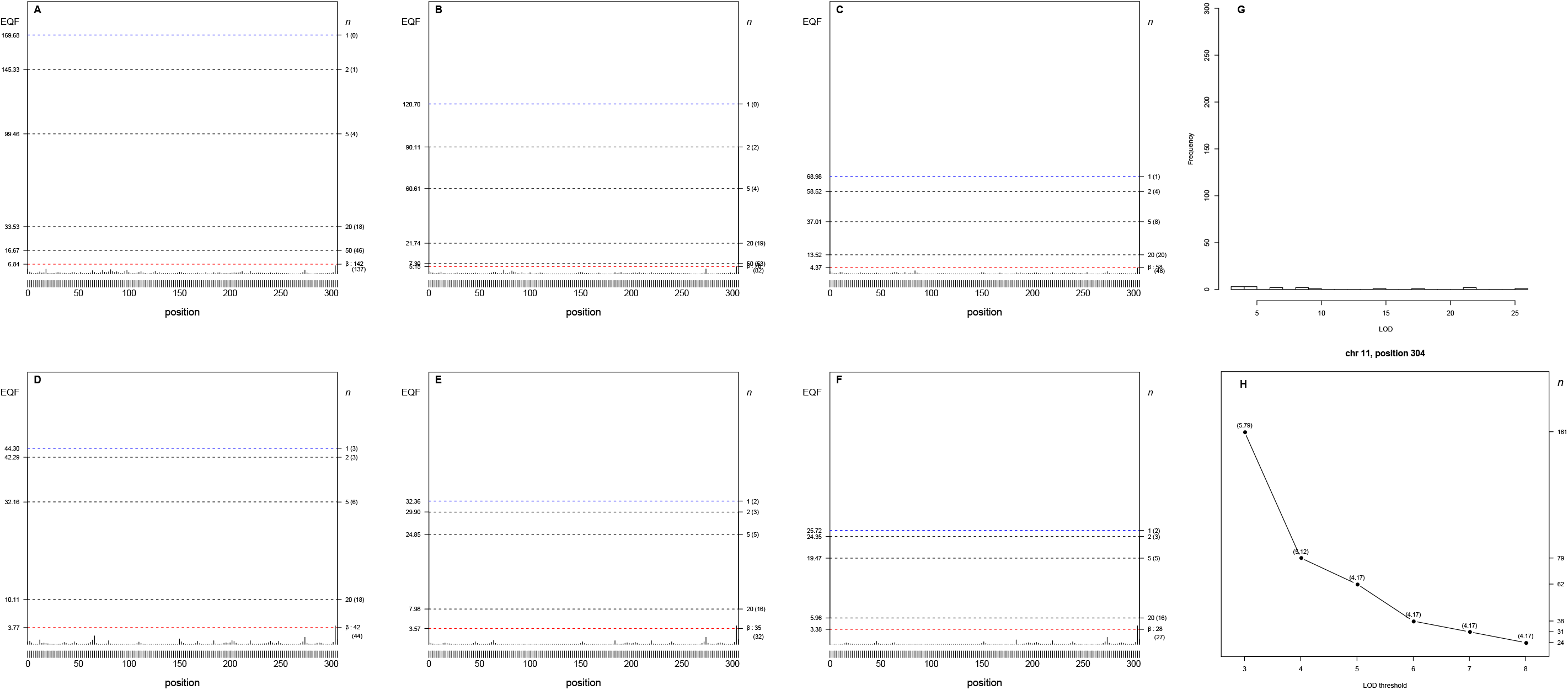
Panels (A-H) The 3- to 8-LOD EQF architectures of the 11^th^ chromosome and the γ_*n*,0.05_ EQF thresholds at GWER of 5%. The left axis denotes the values of EQF, and the right axis denotes the values of n. (A-F) It shows the one peak at the bin [11,304] is significant as the hotspots under the different γ_*n*,0.05_ EQF thresholds in the 3- to 8-LOD EQF architectures. The number in the bracket is the number of detected hotspots. (G) The distribution of LOD scores > 3 for the QTL at the bin [11,304]. (H) The top γ_*n*,0.05_ profile at the bin [11,304] shows that the values of n have a flat pattern varying from 161 to 24 across the 3- to 8-LOD EQF architectures. The number in the bracket is the EQF value of the bin.

**Figure S3-16.**
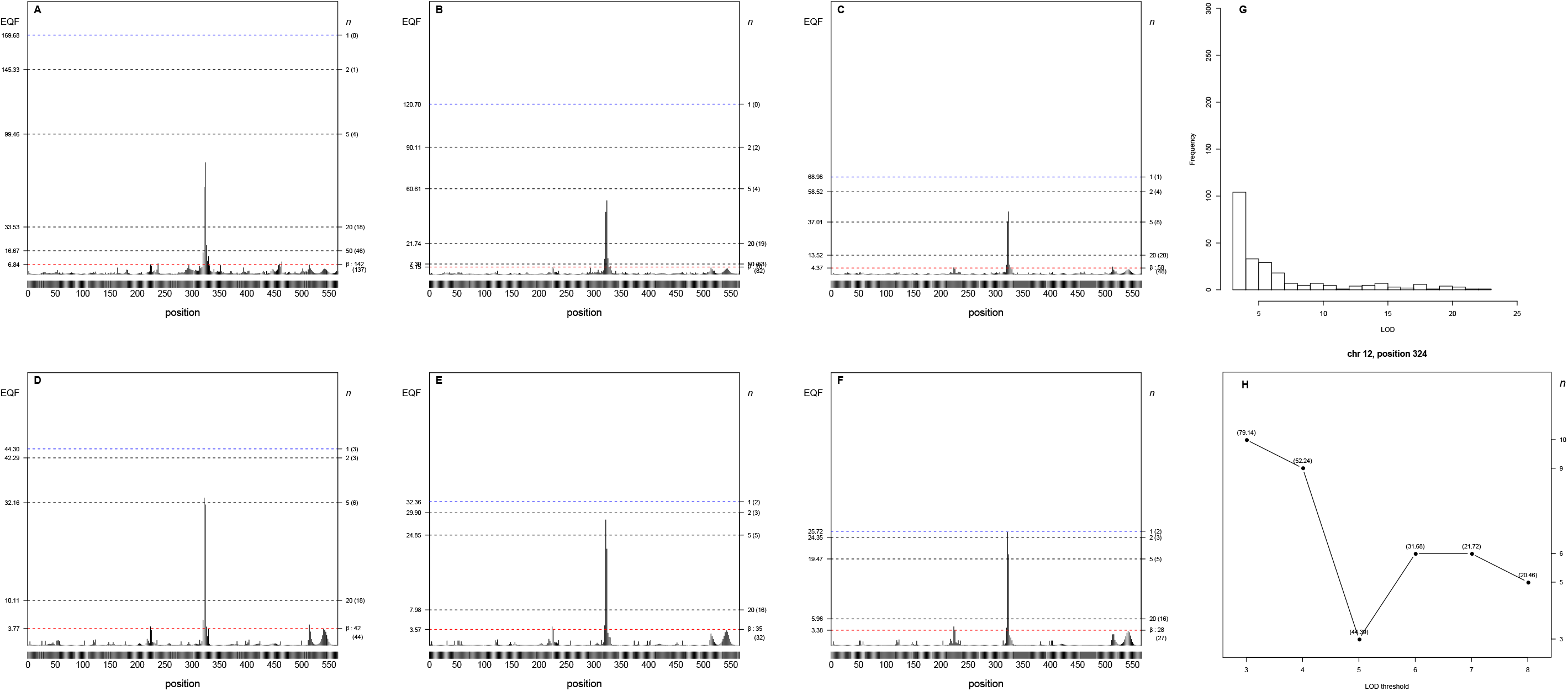
Panels (A-H) The 3- to 8-LOD EQF architectures of the 12^th^ chromosome and the γ_*n*,0.05_ EQF thresholds at GWER of 5%. The left axis denotes the values of EQF, and the right axis denotes the values of n. (A-F) It shows the one peak at the bin [12,324] is significant as the hotspots under the different γ_*n*,0.05_ EQF thresholds in the 3- to 8-LOD EQF architectures. The number in the bracket is the number of detected hotspots. (G) The distribution of LOD scores > 3 for the QTL at the bin [12,324]. (H) The top γ_*n*,0.05_ profile at the bin [12,324] shows that the values of n have a flat pattern varying from 10 to 3 across the 3- to 8-LOD EQF architectures. The number in the bracket is the EQF value of the bin.

**Figure S3-17.**
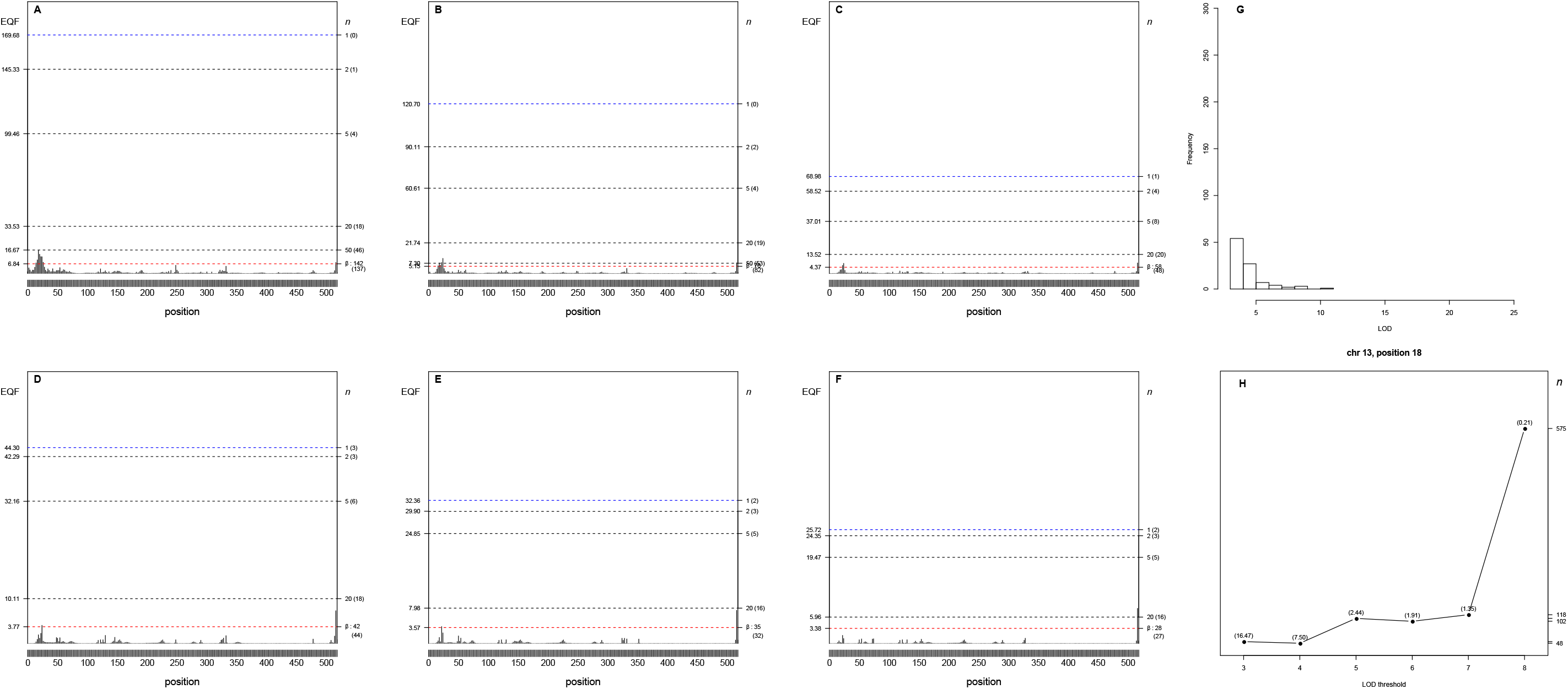
Panels (A-H) The 3- to 8-LOD EQF architectures of the 13^th^ chromosome and the γ_*n*,0.05_ EQF thresholds at GWER of 5%. The left axis denotes the values of EQF, and the right axis denotes the values of n. (A-F) It shows the one peak at the bin [13,18] is significant as the hotspots under the different γ_*n*,0.05_ EQF thresholds in the 3- to 8-LOD EQF architectures. The number in the bracket is the number of detected hotspots. (G) The distribution of LOD scores > 3 for the QTL at the bin [13,18]. (H) The top γ_*n*,0.05_ profile at the bin [13,18] shows that the values of n have a flat pattern varying from 48 to 575 across the 3- to 8-LOD EQF architectures. The number in the bracket is the EQF value of the bin.

**Figure S3-18.**
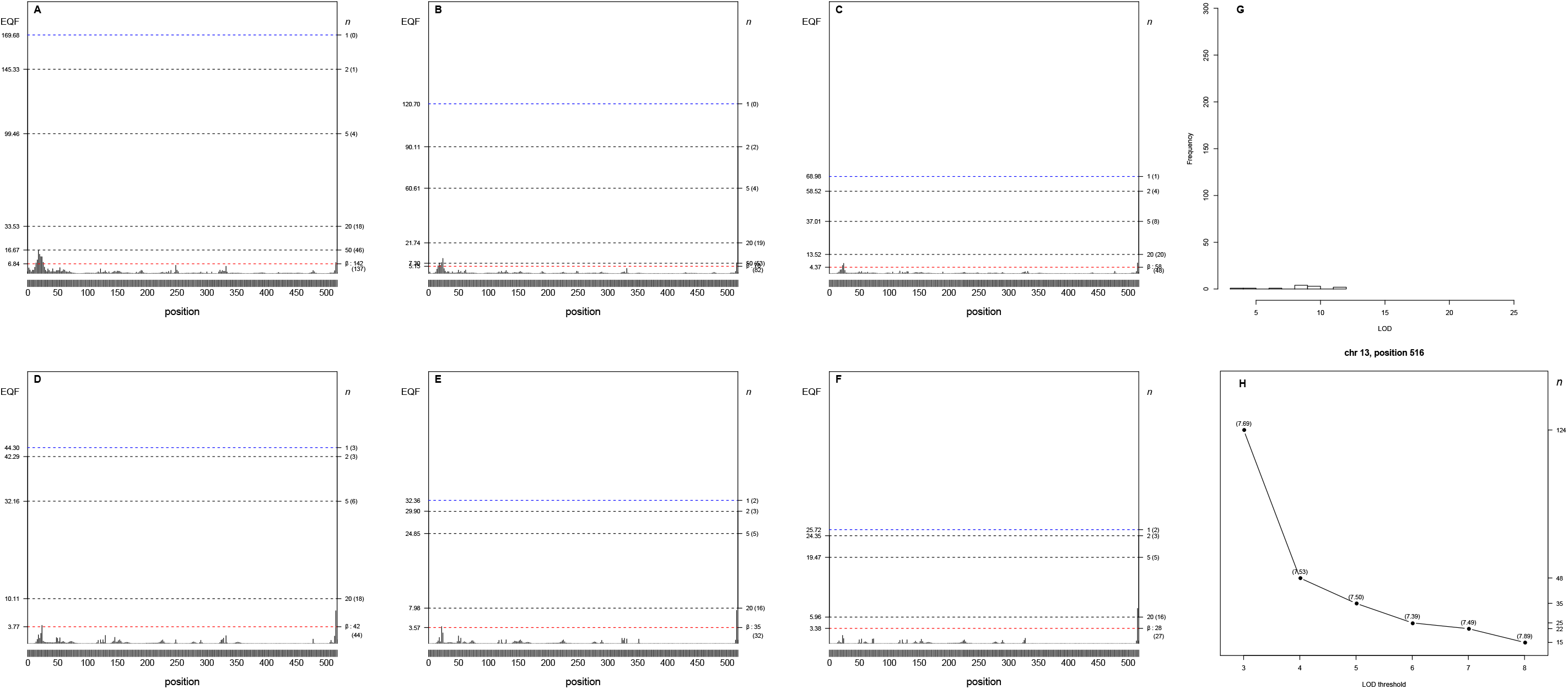
Panels (A-H) The 3- to 8-LOD EQF architectures of the 13^th^ chromosome and the γ_*n*,0.05_ EQF thresholds at GWER of 5%. The left axis denotes the values of EQF, and the right axis denotes the values of n. (A-F) It shows the one peak at the bin [13,516] is significant as the hotspots under the different γ_*n*,0.05_ EQF thresholds in the 3- to 8-LOD EQF architectures. The number in the bracket is the number of detected hotspots. (G) The distribution of LOD scores > 3 for the QTL at the bin [13,516]. (H) The top γ_*n*,0.05_ profile at the bin [13,516] shows that the values of n have a flat pattern varying from 124 to 15 across the 3- to 8-LOD EQF architectures. The number in the bracket is the EQF value of the bin.

**Figure S3-19.**
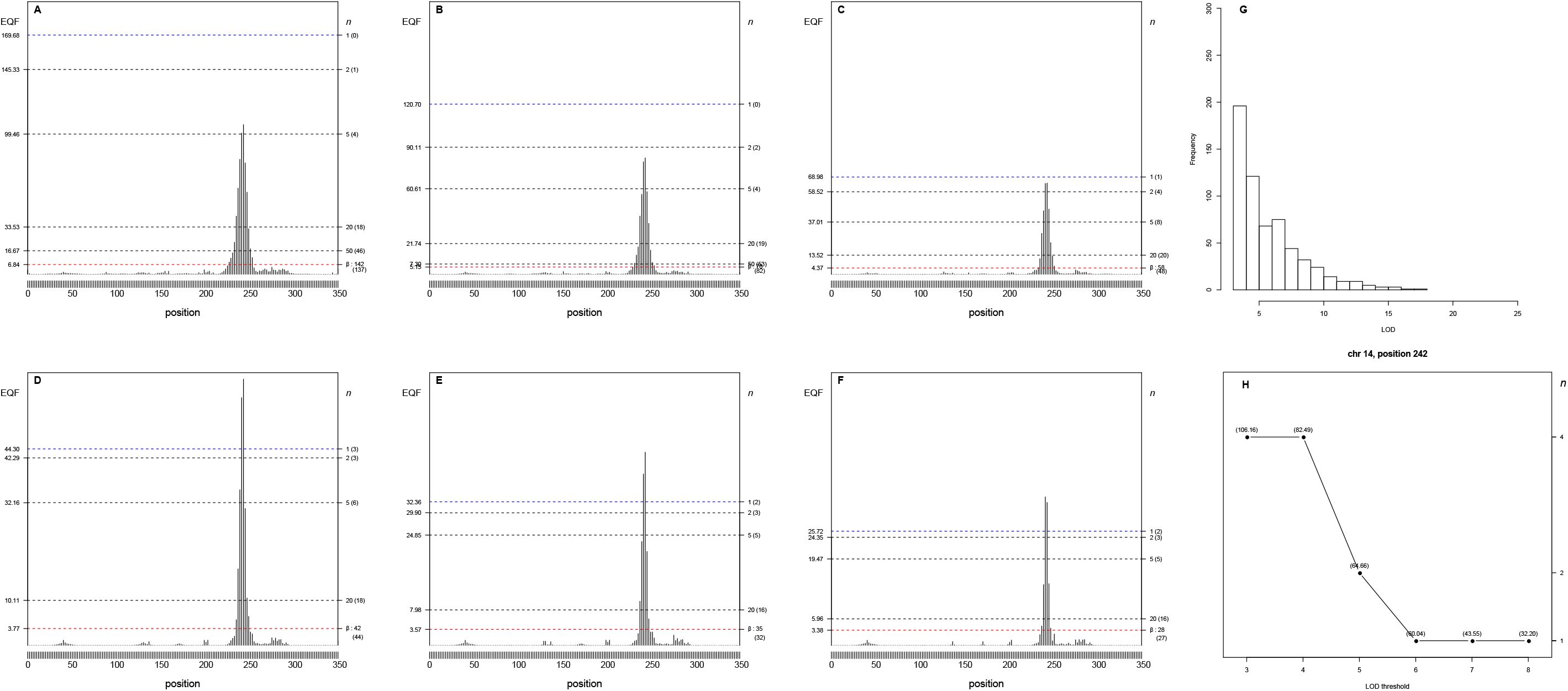
Panels (A-H) The 3- to 8-LOD EQF architectures of the 14^th^ chromosome and the γ_*n*,0.05_ EQF thresholds at GWER of 5%. The left axis denotes the values of EQF, and the right axis denotes the values of n. (A-F) It shows the one peak at the bin [14,242] is significant as the hotspots under the different γ_*n*,0.05_ EQF thresholds in the 3- to 8-LOD EQF architectures. The number in the bracket is the number of detected hotspots. (G) The distribution of LOD scores > 3 for the QTL at the bin [14,242]. (H) The top γ_*n*,0.05_ profile at the bin [14,242] shows that the values of n have a flat pattern varying from 4 to 1 across the 3- to 8-LOD EQF architectures. The number in the bracket is the EQF value of the bin.

**Figure S3-20.**
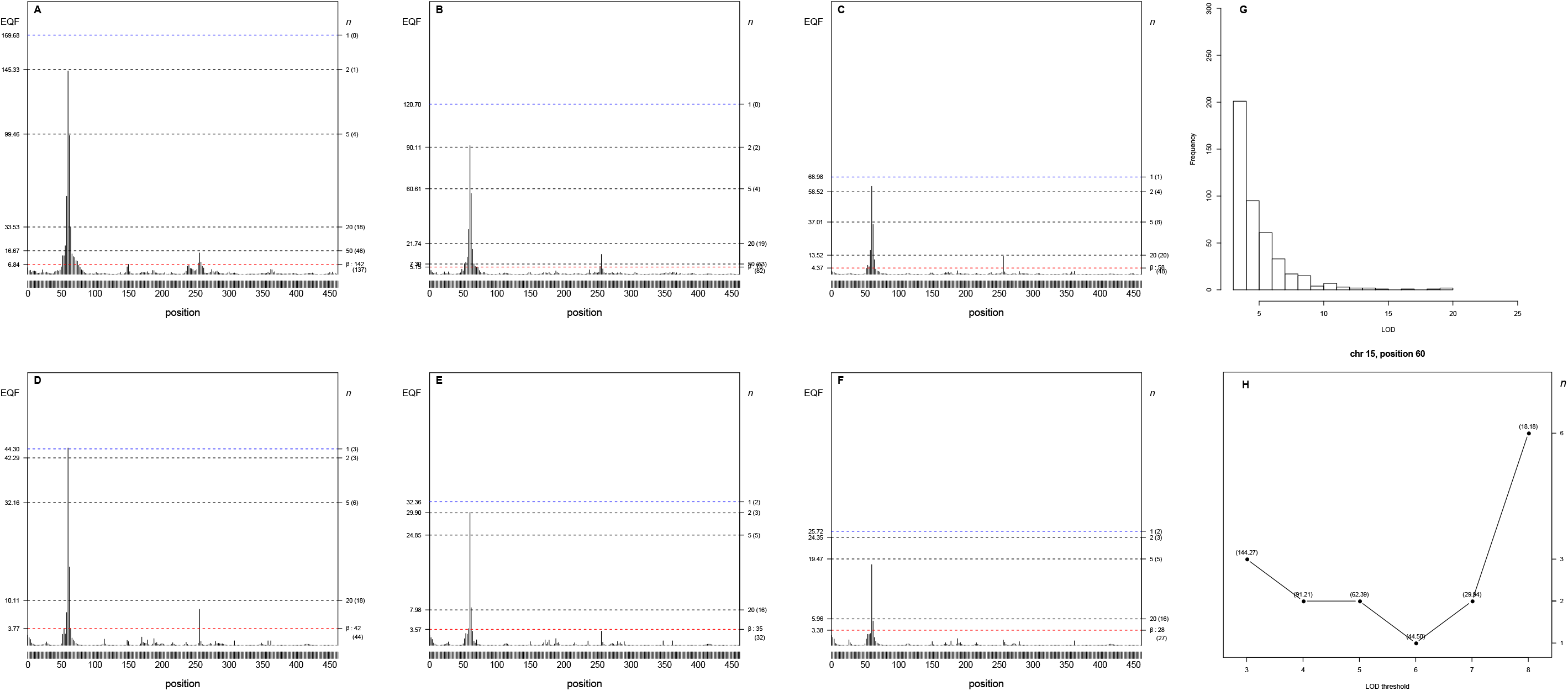
Panels (A-H) The 3- to 8-LOD EQF architectures of the 15^th^ chromosome and the γ_*n*,0.05_ EQF thresholds at GWER of 5%. The left axis denotes the values of EQF, and the right axis denotes the values of n. (A-F) It shows the one peak at the bin [15,60] is significant as the hotspots under the different γ_*n*,0.05_ EQF thresholds in the 3- to 8-LOD EQF architectures. The number in the bracket is the number of detected hotspots. (G) The distribution of LOD scores > 3 for the QTL at the bin [15,60]. (H) The top γ_*n*,0.05_ profile at the bin [15,60] shows that the values of n have a flat pattern varying from 1 to 6 across the 3- to 8-LOD EQF architectures. The number in the bracket is the EQF value of the bin.

**Figure S3-21.**
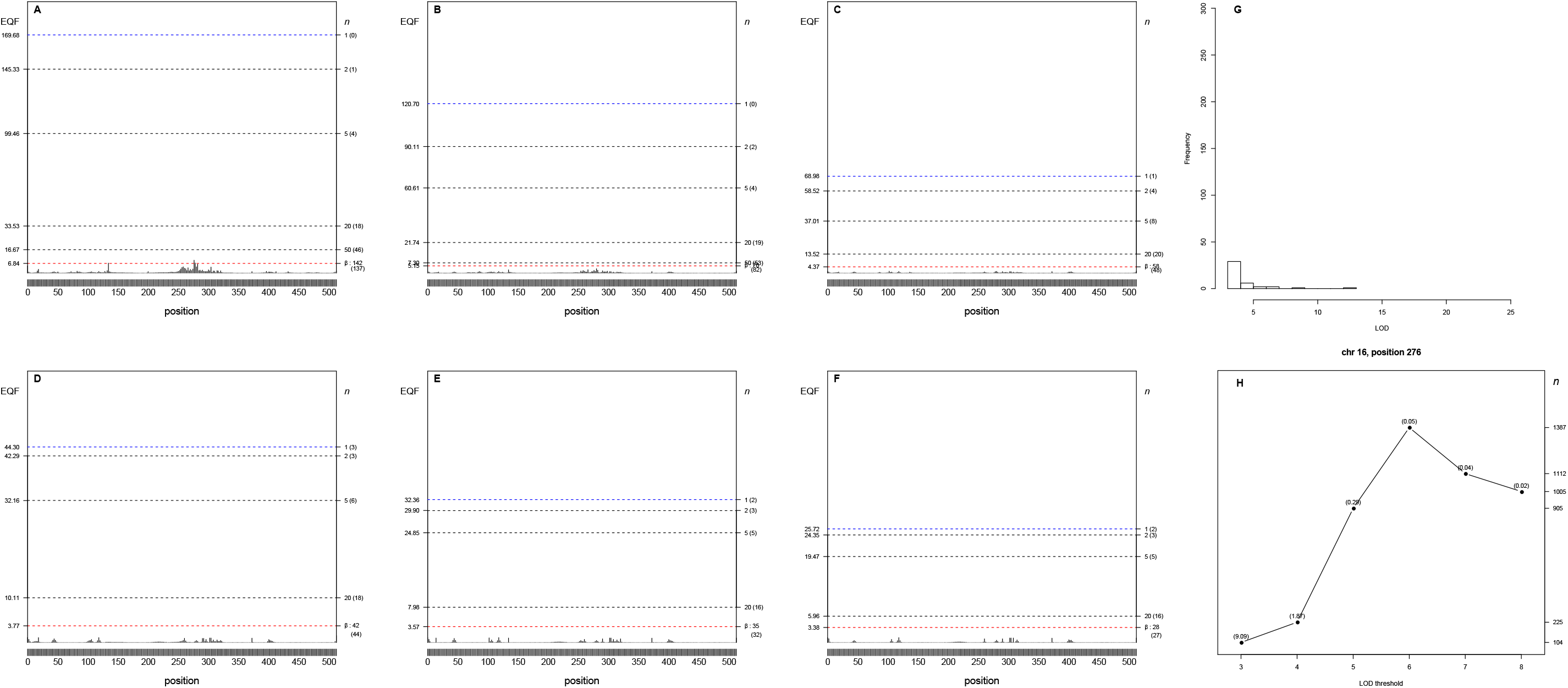
Panels (A-H) The 3- to 8-LOD EQF architectures of the 16^th^ chromosome and the γ_*n*,0.05_ EQF thresholds at GWER of 5%. The left axis denotes the values of EQF, and the right axis denotes the values of n. (A-F) It shows the one peak at the bin [16,276] is significant as the hotspots under the different γ_*n*,0.05_ EQF thresholds in the 3- to 8-LOD EQF architectures. The number in the bracket is the number of detected hotspots. (G) The distribution of LOD scores > 3 for the QTL at the bin [16,276]. (H) The top γ_*n*,0.05_ profile at the bin [16,276] shows that the values of n have a flat pattern varying from 104 to 1378 across the 3- to 8-LOD EQF architectures. The number in the bracket is the EQF value of the bin.

